# Mitochondrial organization in the developing proximal tubule is controlled by LRRK2

**DOI:** 10.1101/2025.09.02.673829

**Authors:** Mohsina Anjum Khan, Kyle Bond, Elyse Grilli, Daniel Cameron, Liyang Zhao, Sunder Sims-Lucas, Andrew P. McMahon, Thomas J. Carroll, Leif Oxburgh

**Author notes:** Division of Biology and Biological Engineering, California Institute of Technology, Pasadena, CA 91125.

## Abstract

The proximal tubule of the nephron resorbs water, amino acids and glucose, and its energy demands are high. Formation of the cellular machinery for energy production is an essential step in proximal tubule epithelial cell (PTC) differentiation, and this report focuses on how mitochondria in nascent PTCs are redistributed from their initial apical position to their ultimate basolateral location. We found that mitochondria move from the apical to basolateral side of the PTC coincident with the initiation of lumen flow and that proximal tubules deficient in filtration (aglomerular mice and kidney organoids) maintain their mitochondria in the apical position, indicating that flow is necessary and sufficient for localization. Further, we show that mitochondrial localization depends on the activity of LRRK2 *in vitro* and *in vivo*. Modeling the effect of fluid flow on PTCs demonstrates that LRRK2 activity is regulated by fluid shear stress, providing an explanation for how onset of flow in the newly differentiated proximal tubule may trigger the apical-to-basolateral dissemination of mitochondria that forms the template for subsequent PTC maturation. Our findings indicate that mitochondrial redistribution is one component of a cellular program in the nascent PTC that drives function and that this process is regulated by flow.

## Introduction

The mammalian kidney provides multiple essential physiological functions, including waste excretion, electrolyte balance, pH balance and control of red blood cell production. This organ can be considered an integral component of the cardiovascular system and receives 15-20% of the cardiac output. As blood passes through the glomerular capillary, water and molecules smaller than the approximately 50 kD cut-off of the glomerular filtration barrier pass through to the nephron. In a human being, the daily quantity of filtrate that passes into the nephron is approximately 160 liters and thus reclamation of water, glucose and amino acids that are not destined for excretion is an absolute necessity for survival. The majority of this reclamation occurs in the nephron segment into which the glomerular filtrate first empties; the proximal tubule (PT). The large volume of resorption required of this nephron segment results in an extreme energy demand, and the PT cell (PTC) is estimated to be the second most energy demanding cell in the human body after the cardiomyocyte. Energy to drive the physiological processes of the PTCs is derived largely from fatty acid oxidation, and thus mitochondrial mass and efficiency are critical determinants of PTC function (Bhargava and Schnellmann, 2017). Several congenital mitochondrial disorders manifest themselves in the kidney (Finsterer and Scorza, 2017), highlighting the critical relationship between mitochondrial function and kidney health.

The functional capacity of mitochondria is determined by their abundance but also by their structure. Mitochondrial biogenesis is controlled by concerted activation of nuclear and mitochondrial genetic programs. While 13 components of the electron transport chain are encoded by the mitochondrial genome, the vast majority of mitochondrial proteins are encoded by the nuclear genome. A family of transcription factors including PPARGC1A (PGC1α), PPARGC1B (PGC1β), and PPRC1 (PRC) acts as master regulator and coordinates synthesis of structural components, metabolic components and the mitochondrial genome (Dorn et al., 2015). Structurally, the balance between fission and fusion of mitochondria determines the proportion of mitochondria that form long, contiguous structures that are energetically efficient. Fission is controlled by the GTPase DRP1, while fusion is controlled by the GTPases mitofusin 1 and 2, and Opa1 (Mishra and Chan, 2014). In actively proliferating cells, fission ensures that mitochondria can be appropriately distributed in daughter cells, while fusion ensures mitochondrial efficiency for the synthetic phase. The subcellular localization of mitochondria is an important feature of differentiated, post-mitotic cells, particularly in cell types where energy-dependent processes are compartmentalized. The machinery for active subcellular transport and localization of mitochondria has been most actively studied in neurons, where mitochondria are transported along axons by physical association of a molecular complex attached to their outer membrane that includes TRAK1/2, RHOT1/2, kinesin and dynein with the microtubule network (Panchal and Tiwari, 2021). Electron microscopy studies have shown that the subcellular localization of mitochondria in kidney epithelial cells is stereotypical, but how mitochondria reach their ultimate destination at the basal aspect of the cell where they are closely associated with the energy-demanding Na-K ATPases has not been reported.

Research over the past decade has refined methods to generate PTs from pluripotent stem cells in vitro, providing new and exciting possibilities to study human kidney biology in three dimensional organoid systems (Morizane et al., 2015; Takasato et al., 2015). Genetic studies support the interpretation that formation of nephrons in human kidney organoids recapitulates the developmental process genetically (Combes et al., 2019; Howden et al., 2019), opening opportunities to study the basic cell biology of PT formation in vitro.

In this study, we asked how mitochondria that are located apically in the nascent and immature PTC are redistributed throughout the cell to reach their ultimate basolateral location. Analysis of dextran uptake by PTCs in embryos and neonates suggests that lumen flow triggers apical-to-basolateral redistribution of mitochondria, a finding that is corroborated by studies of mice that are genetically deficient in glomeruli. Membrane potential analyses suggests that the apical to basolateral transition of mitochondria associates with increased energy output. Using human kidney organoids and neonatal mice, we show that mitochondria are actively apically sequestered in nascent PTCs prior to onset of filtration and that this process is dependent on the multifunctional kinase LRRK2. Modeling fluid flow on PTCs demonstrates that LRRK2 activity is regulated by fluid shear stress, explaining how onset of filtration in the newly differentiated PT can trigger the apical-to-basolateral dissemination of mitochondria characteristic of the mature PTC.

## Results

### Mitochondrial mass increases and mitochondria localize basolaterally during PT maturation

To define mitochondrial organization during early differentiation steps of the PT, we analyzed kidney tissue from 2 second trimester (week 18-20) human fetuses for molecular markers of PTs and mitochondria (Fig. 1A-H, S1A-C). The process of new nephron addition in the fetal kidney is radial, with specification of new nephrons occurring in the most cortical nephrogenic zone (NZ). Nascent nephrons first form PTs immediately below the NZ, and further growth and maturation of PTs takes place in the deeper cortical layers of the developing kidney. Co-staining with PT molecular markers (HNF4A and LTL) and an antibody against the mitochondrial outer membrane protein TOMM20 shows very early PT precursors with small lumens (PT-early) directly adjacent to the NZ in the outer cortex of the kidneys, and larger PTs with distended lumens (PT-diff) located deep in the cortex (Fig. 1A).

**Figure 1:**
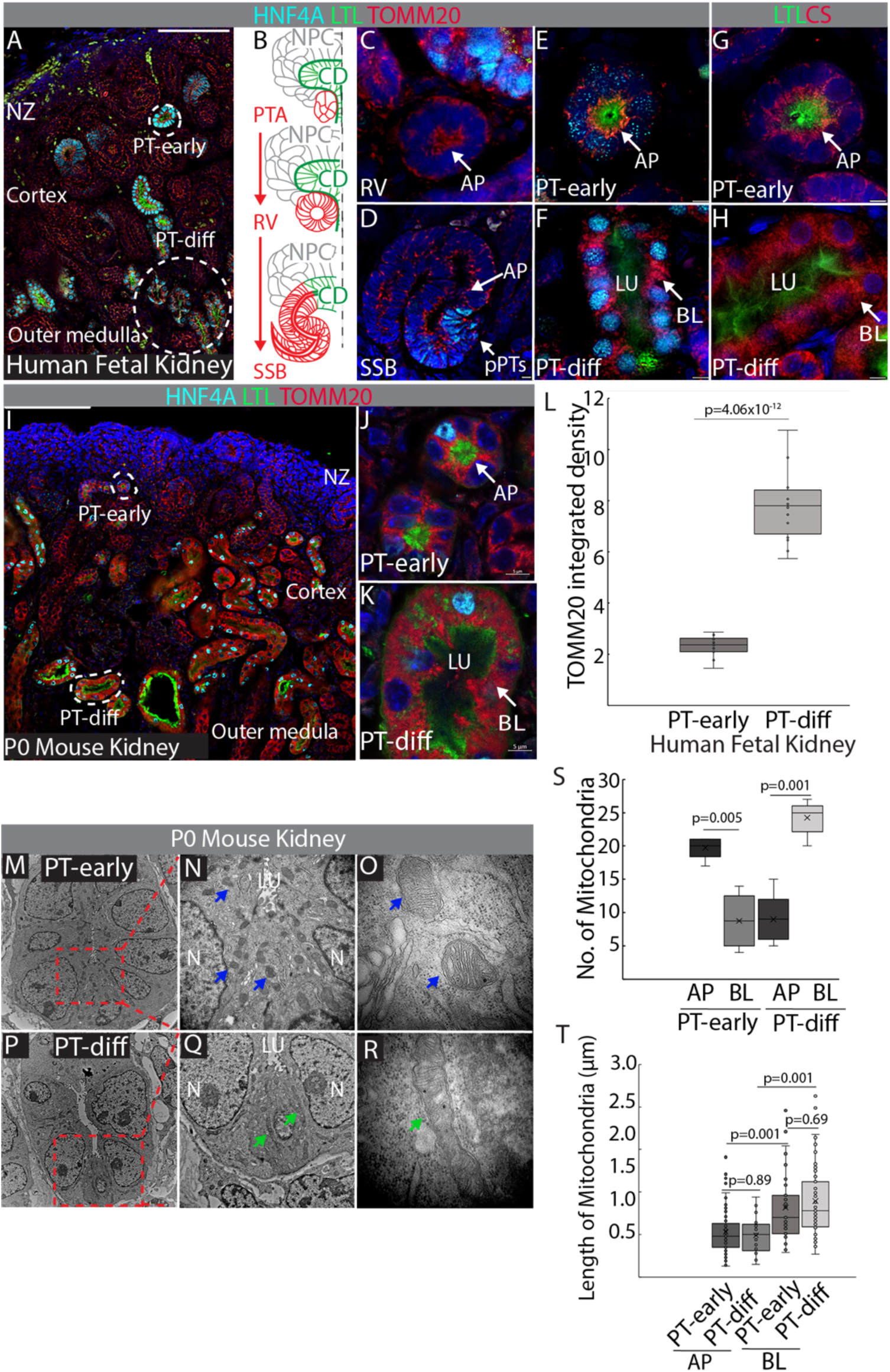
Mitochondrial mass and localization changes during PT differentiation. (A-H) Kidney tissue from 18-20 week human fetus stained with (A-F) PT markers (HNF4A, cyan; LTL, green) and mitochondrial marker (TOMM20, red). (A) Widefield image showing nephrogenic zone, cortex and medulla and location of nephrogenic zone-adjacent HNF4A+/LTL+ PTs (PT-early) with narrow lumens and deep cortical HNF4A+/LTL+ PTs (PT-diff) with distended lumens. (B) Schematic of nephron differentiation. (C) Renal vesicle. (D) S-shaped body with specification of HNF4A+ presumptive PT. (E) PT-early with mainly apical mitochondria. (F) PT-diff with distended lumen and basolateral mitochondria. (G) PT-early and (H) PT-diff stained with the mitochondrial marker citrate synthase (CS, red) and the PT marker LTL (green). (I-K) P0 mouse kidney tissue section. (I) Widefield image showing nephrogenic zone, cortex and medulla and location of nephrogenic zone-adjacent HNF4A+/LTL+ PTs (PT-early) with narrow lumens and deep cortical HNF4A+/LTL+ PTs (PT-diff) with distended lumens. (J) PT-early with mainly apical mitochondria. (K) PT-diff with distended lumen and basolateral mitochondria. (L) Comparison of mean integrated density of TOMM20/cell in human fetal PTs (n=16) with apical mitochondria versus PTs with basolateral mitochondria (n=16). Results are aggregated from 2 different human fetal samples. Data are shown as box-and-whisker with mean (middle line) and minimum–maximum values (whiskers); two-tailed Student’s *t*-test. (M-R) Transmission electron micrographs of P0 mouse kidney showing (M-O) PT-early with mainly apically located rounded mitochondria (blue arrows). (P-R) PT-diff with distended lumen and elongated basolateral mitochondria (green arrows). (S) Comparison of apical to basolateral mitochondrial number in PT-early (n=3) versus PT-diff (n=3). (T) Comparison of length of apical and basolateral mitochondria in PT-early (n=3) and PT-diff (n=3). Data (S and T) is represented as box-and-whisker with mean (middle line) and minimum– maximum values (whiskers); one way ANOVA with post-hoc Tukey HSD test. **Abbreviations:** AP, apical; BL, basolateral; CD, collecting duct; LU, lumen; NPC, nephron progenitor cell; N, nucleus; NZ, nephrogenic zone; pPT, presumptive proximal tubule; PT, proximal tubule; PTA, pretubular aggregate; RV, renal vesicle; SSB, s-shaped body. All immunofluorescence images are counterstained with the nuclear stain DAPI (blue). Scale bars in A & I 100 µm, all other panels 5 µm. The schematic in panel 1B was adapted from (Yang et al., 2013) with permission.

Within the NZ, induced progenitor cells form pretubular aggregates that epithelialize to spheroids of epithelial cells known as renal vesicles (RVs), that are patterned to S-shaped bodies (SSBs) as they fuse their lumens to the collecting duct (Fig. 1B). Proximal-distal patterning of the nephron tubule occurs during the RV to SSB transition and the transcription factor HNF4A is one of the earliest markers of PT identity. Binding of lotus tetragonolobus lectin (LTL) increases as the PT lumen matures and develops an apical glycocalyx. At the RV stage, mitochondria are largely located at the apex of epithelial cells, with limited basolateral distribution (Fig. 1C). In the SSB, both apical and basolateral distribution is seen, but in the HNF4A+/LTL-presumptive PT segment, mitochondria are almost exclusively apically located (Fig. 1D). This pattern of mitochondrial localization is maintained in early HNF4A+/LTL+ PTs found immediately adjacent to the NZ (Fig. 1E). However, HNF4A+/LTL+ PTs in the deep cortex with distended lumens display dramatically increased mitochondrial abundance as well as distribution throughout the cytoplasm (Fig. 1F). To verify that our observations based on TOMM20 reflect mitochondrial localization rather than simply accumulation of this specific protein, we repeated our analysis using an antibody that recognizes citrate synthase (CS), which is located within the mitochondrial matrix. We found an identical staining pattern, with mitochondria in early HNF4A+/LTL+ PTs located apically (Fig. 1G), while more medullary PTs showed distribution throughout the cytoplasm (Fig. 1H). To ascertain if the redistribution of apical mitochondria during PT differentiation is conserved between mouse and human, we conducted the same immunostaining analysis on neonatal mouse kidneys. Active nephrogenesis continues until day 3 after birth in the mouse, and the stratification of PT differentiation stages from NZ to medulla is very pronounced (Fig. 1I). Similar to the human fetal kidney, early HNF4A+/LTL+ PTs beneath the NZ in neonatal mouse kidneys had primarily apical mitochondria (Fig. 1J), while PTs in the deep cortex had drastically increased mitochondrial mass distributed throughout the cytoplasm (Fig. 1K).

Quantitative comparison of mitochondrial abundance per cell based on integrated density of fluorescent signal showed an approximately 4-fold increase in mitochondrial abundance between PTs with apical mitochondria and PTs with mitochondria distributed throughout the cytoplasm in human fetal kidneys (Fig. 1L), demonstrating that both biogenesis and subcellular localization of mitochondria change as PTs mature following their initial specification.

To confirm our observations regarding differences in mitochondrial localization between PT-early and PT-diff, we performed transmission electron microscopy on P0 kidneys (Fig. 1M-T). PT-early display abundant rounded mitochondria apically located between the nuclei and the tubule lumen (Fig. 1M,N). These were identified as mitochondria based on their outer bilayer and cristae (Fig. 1O). Basolateral mitochondria can only be located sporadically in PT-early. In PT-diff, apical mitochondria are rare, and extended mitochondria can be seen adjacent to nuclei (Fig. 1P,Q). Higher magnification imaging reveals that these mitochondria appear many times longer than the apical mitochondria found in PT-early, and that they have extensive cristae formation (Fig. 1R). Differences in both mitochondrial number and mitochondrial length are supported by quantitative analysis of cross sections of 3 PT-early versus 3 PT-diff from different kidneys. Figure 1S shows comparison of apical mitochondria located between the lumen and the nucleus versus basolateral mitochondria located adjacent to the nucleus down to the level of the basement membrane in PT-early versus PT-diff. Figure 1T shows comparison of length of apical versus basolateral mitochondria in PT-early versus PT-diff. While there is a significant size difference between apically and basolaterally located mitochondria in both PT-early and PT-diff, there is no detectable difference in size of either apical or basolateral mitochondria between PT-early and PT-diff.

We were curious to understand if the apical localization of mitochondria in PT-early could be unique to this tubule segment, so we repeated our mitochondrial staining together with the distal tubule marker lycopersicon esculentum lectin (LEL) (Fig. S1D-K). Widefield imaging shows extensive differentiation of LEL+ tubules in the neonatal mouse kidney (Fig. S1D), allowing us to select tubule cross-sections adjacent to PT-early and PT-diff for analysis. Compared with PT-early, adjacent LEL+ tubules (designated DT-1 (Fig. S1E)) display basolateral and more abundant mitochondria (Fig. S1F-G’). Compared with PT-diff, adjacent LEL+ tubules (designated DT-2 (Fig. S1H)) display basolateral mitochondria that are less abundant (Fig. S1I-J’). Quantification supports these observations (Fig. S1K), suggesting that the pronounced apical-to-basolateral re-location of mitochondria from PT-early to PT-diff may be unique to the proximal segment of the nascent nephron.

### Distribution of mitochondria from their apical localization during the differentiation process associates with increased mitochondrial polarization

While it can be inferred from several genetic and organ culture studies that there is an important role for mitochondrial metabolism in the differentiation of nephron progenitor cells to epithelium (Cargill et al., 2019; Ishii et al., 2021; Liu et al., 2017), there is a paucity of spatial data on mitochondrial activity in the developing kidney. To understand the significance of the changes in mitochondrial mass and localization identified in this study, we labeled P0 mouse kidney tissue with tetramethylrhodamine methyl ester (TMRM), which accumulates and fluoresces in mitochondria proportionally to membrane potential (Ehrenberg et al., 1988). TMRM has previously been used to measure PT mitochondrial activity in the adult rat using *in vivo* multiphoton microscopy. In this report, peak fluorescence within PTs was reached within 5 minutes after intravenous injection and maintained for 45 minutes at over 80% of the peak signal strength (Hall et al., 2013). Based on these metrics, we injected P0 mice intravenously with TMRM, sacrificed them after 30 minutes, harvested the kidneys and imaged them within 5 minutes of sacrifice to stay within the 45 minutes time window during which there is minimal signal decay (Fig. 2A,S2A). One important observation from intravital imaging is that TMRM is mainly taken up basally by the cells of the tubule epithelium (Hall et al., 2013), which is consistent with uptake directly from the peritubular vasculature. We reasoned that the paucity of signal seen in the NZs of P0 kidneys that were TMRM loaded through intravenous injection (Fig. S2A) might be due to the lack of perfusion in the NZ of the developing kidney (Rymer et al., 2014). To answer this question, we bisected freshly harvested kidneys and immersed them in TMRM for 30 minutes, thereafter imaging within the same timeframe (Fig. S2B). Comparing intravenously loaded and immersed kidneys, we observed an overlap in signal with the exception of the NZ, where immersed kidneys showed strong TMRM loading in sparsely distributed epithelial tubules (compare Fig. S2A’ with S2B’). While we performed TMRM loading studies using both intravenous injection and immersion, data presented henceforth is from experiments based on immersion.

**Figure 2:**
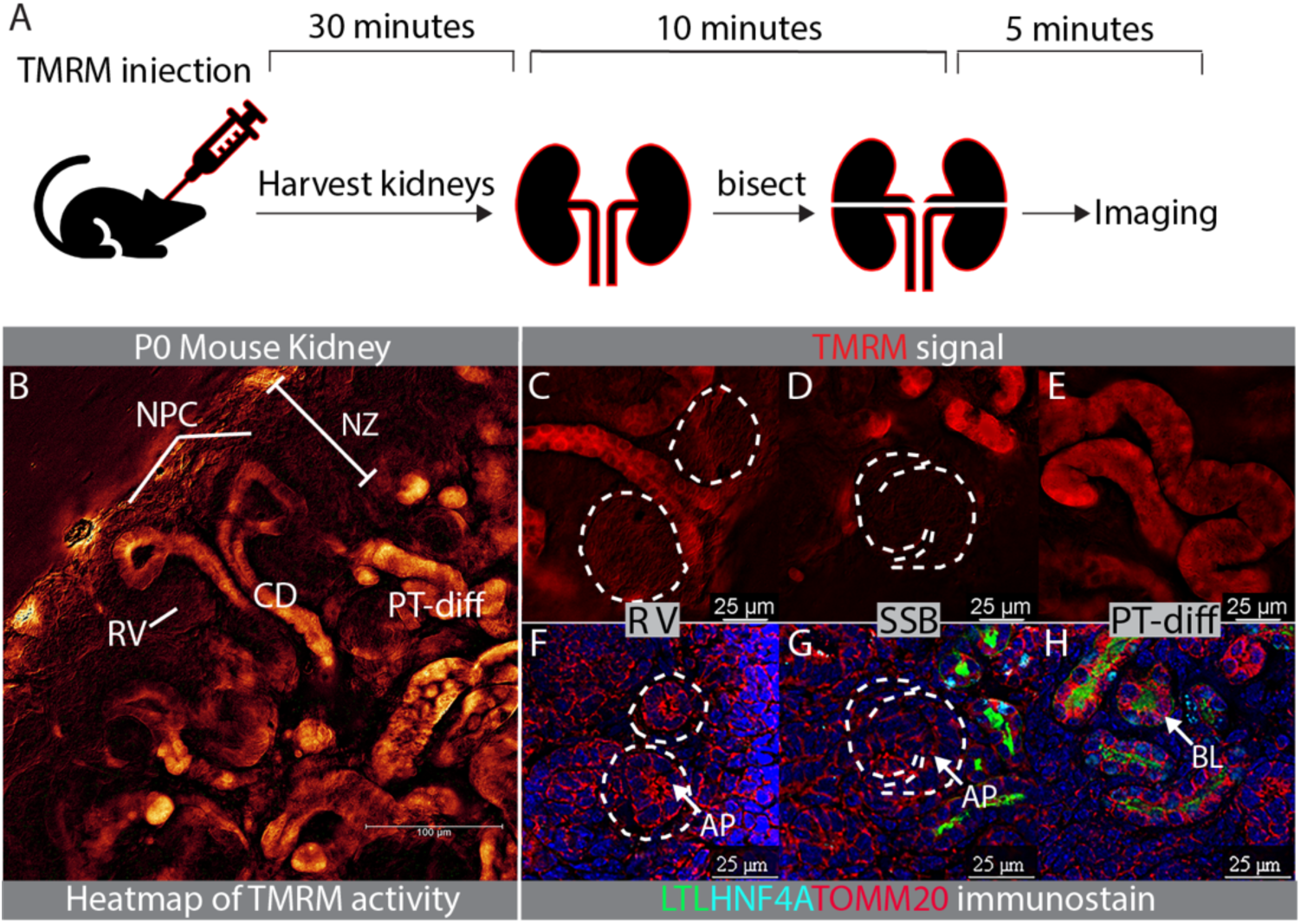
Correlation between mitochondrial activity and mitochondrial localization. (A) Schematic of approach for visualization of TMRM fluorescence in kidney following intravenous injection. (B) Representative CCCP-corrected heatmap from 8 individual measurements of red fluorescence in kidneys immersed in TMRM for 30 minutes. (C) TMRM staining in renal vesicles outlined with dashed lines. (D) TMRM staining in S-shaped body outlined with dashed line. (E) TMRM staining in nephron tubule. (F) Mitochondrial localization in renal vesicles outlined with dashed line (G) Mitochondrial localization in S-shaped body outlined with dashed line. (H) Newly formed cortical nephron tubule counterstained with LTL (green) and HNF4A (cyan) to identify PTs. **Abbreviations:** AP, apical; BL, basolateral; CD, collecting duct; NPC, nephron progenitor cell; NZ, nephrogenic zone; PT, proximal tubule; RV, renal vesicle; SSB, s-shaped body. Immunofluorescence images F, G, H were counterstained with the nuclear dye DAPI (blue).

TMRM signal can be strongly influenced by efflux through multi drug resistance (MDR) pumps, and we therefore included conditions in which we immersed kidneys in three different concentrations of the MDR inhibitor verapamil prior to addition of TMRM. As reported for intravital analysis of rat PT (Hall et al., 2013), addition of verapamil does not affect TMRM signal intensity indicating that efflux through MDR pumps is not a significant confounder (Fig. S2C). To gauge the level of tissue autofluorescence in the assay, we pretreated kidneys with three doses of the mitochondrial membrane depolarizer CCCP (Fig. S2D). We subtracted the background in CCCP-treated sections from the signal in sections immersed only in TMRM to generate a heatmap of TMRM fluorescence as a proxy measurement of mitochondrial membrane polarization in the P0 mouse kidney (Fig. 2B). Heatmaps were generated by imaging kidneys from 8 different individuals, the same pattern was seen in each and a representative heatmap is shown. Using this approach, we found that the NZ is largely devoid of signal with the exception of the collecting ducts (Fig. 2B,S2B’). Renal vesicles (Fig. 2C) and S-shaped bodies (Fig. 2D) show little or no signal, while nascent tubules associated with primitive glomeruli show clear signal (Fig. 2E), indicating that mitochondrial polarization increases with differentiation of the tubule epithelium.

To determine the relationship between apical localization of mitochondria and polarization we compared TMRM activity and TOMM20 signal between the renal vesicle (Fig. 2C vs F), S-shaped tubule precursors (Fig. 2D vs G) and tubule segments connected to a glomerulus (Fig. 2E vs H). Little or no mitochondrial polarization is visible in tubule precursors, while PTs associated with glomeruli display polarization. Based on these observations, we propose that the redistribution of apical mitochondria is associated with a significant increase in mitochondrial polarization and energy production. This correlates well with our electron microscopy observation that mitochondria are small and fragmented apically, but elongated basolaterally, suggesting higher energetic efficiency.

### Onset of lumen flow coincides with basolateral localization of mitochondria

Basolateral localization of mitochondria occurs at an early point in PT differentiation, and one possibility is that it is associated with the onset of lumen flow. To investigate this relationship, we performed dextran injections to identify functional PTCs and compare mitochondrial localization between pre-functional and functional PTCs in nascent nephrons. Individual PTs with lumen flow were labeled by intravenous injection with fluorescently tagged 10 kD dextran, which passes across the glomerular filtration barrier into the lumens of filtering nephrons, where it is taken up apically by PTCs through fluid-phase endocytosis. Endocytosis is activated by fluid shear stress in PTCs (Long et al., 2017), and dextran uptake thus serves as a sensitive marker for onset of flow in the nephron. To determine the appropriate chase period after dextran injection, we performed a series of pulse-chase experiments in P0 mice and found clearance of dextran solution from the tubule lumen and clear apical uptake of dextran in endosomes 30 minutes after intravenous injection (Fig. 3A, data not shown). HNF4A+ PTs with uptake of dextran displayed strictly basolateral mitochondrial localization at P0 while PTs without dextran uptake displayed apical mitochondria (Fig. 3B-F), suggesting that basolateral localization of mitochondria indeed does coincide with onset of lumen flow. These observations in neonatal kidneys suggest that PT-early are HNF4A+/dextran-negative and pre-functional whereas PT-diff are HNF4A+/dextran-positive and functional.

**Figure 3:**
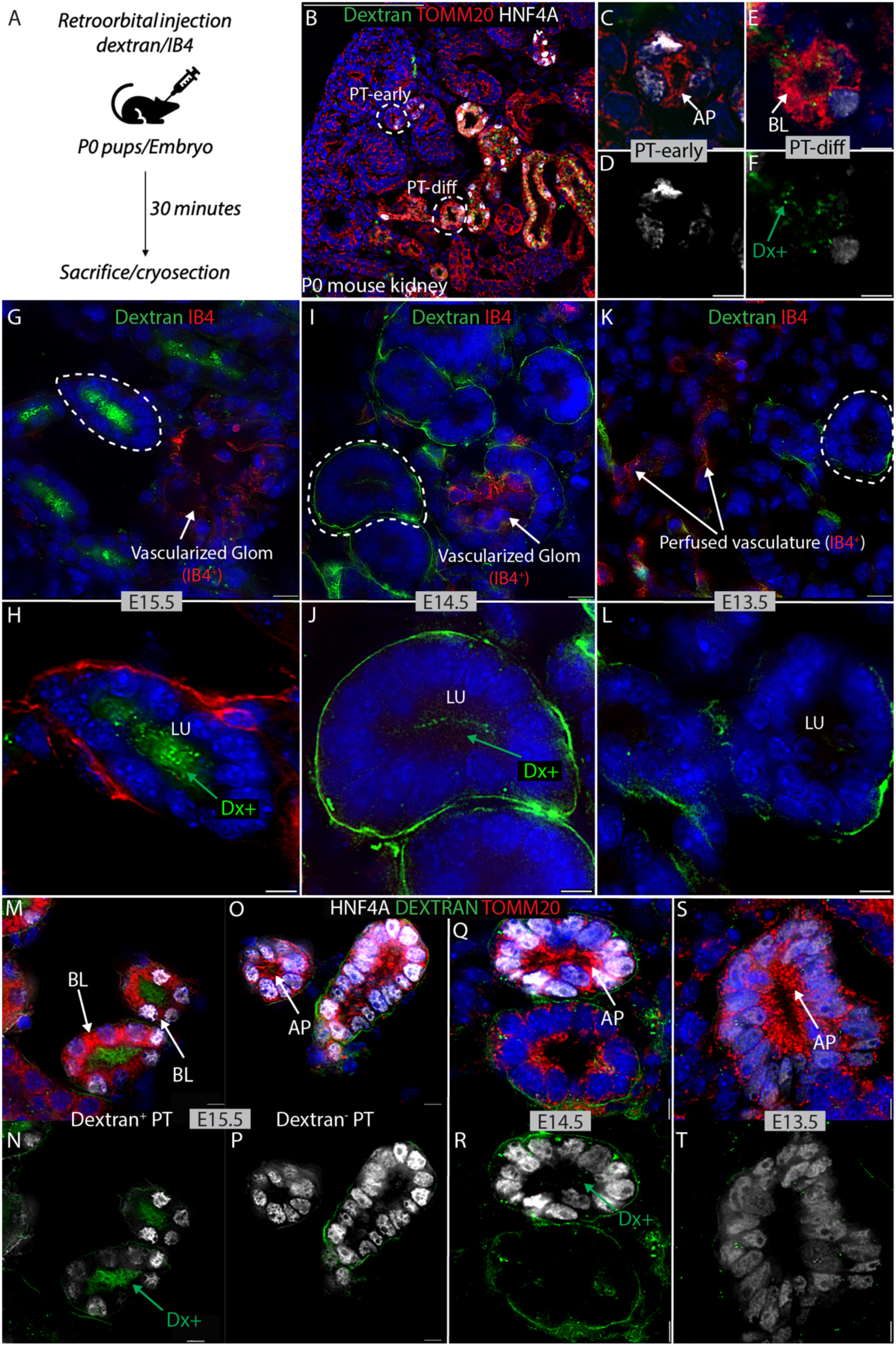
Correlation between mitochondrial localization and onset of function in PTs. (A) Approach for intravital labeling of functional PTs with Alexafluor-488-labeled 10K dextran to distinguish between pre-functional and functional PTs. (B) Widefield image of dextran-labeled (green) P0 mouse kidney, stained with DAPI (nucleus, blue), TOMM20 (mitochondria, red), and HNF4A (PT, white). (C, D) Dextran-negative, pre-functional PTs. (E, F) Dextran-positive, functional PTs. (G, H) E15.5, (I, J) E14.5, (K,L) E13.5 kidneys from embryos intravenously injected with dextran (green) and Alexafluor-647-labeled IB4 (red). DAPI (nucleus, blue), TOMM20 (mitochondria, red), HNF4A (PT, white), dextran (green) labeling of (M, N) E15.5 tubules with apical dextran uptake; (O, P) E15.5 tubules without apical dextran uptake; (Q, R) E14.5 tubules with traces of apical dextran uptake; (S,T) E13.5 tubules without apical dextran uptake. **Abbreviations:** AP, apical; BL, basolateral; Dx, dextran; LU, lumen; PT, proximal tubule. All immunofluorescence images are counterstained with the nuclear stain DAPI (blue). Scale bar in B 100 µm, all other panels 5 µm.

If basolateral localization of mitochondria coincides with onset of lumen flow, then we would expect embryonic nephrons to display apically located mitochondria in PTs at developmental stages prior to onset of lumen flow. To test this, we performed intravenous dextran labeling through retroorbital injection on a series of mouse embryos aged E13.5, E14.5, E15.5, and E17.5. As a tracer to confirm dye distribution to the kidney vasculature, we injected a cocktail of Alexafluor-647-labeled Griffonia simplificata isolectin B4 (IB4) and Alexafluor-488-labeled 10 kD dextran. IB4 labels glycoproteins on the endothelial lumen in the mouse, and we have previously used it as an intravital marker of perfused kidney vasculature (Kumar Gupta et al., 2020). Imaging of the Alexa-488 wavelength in whole embryos 1 minute after injection confirmed distribution of the labeled dextran throughout the vasculature, including the posterior region (Fig. S3A-C). Kidneys were harvested after 30 minutes and cryosectioned. Imaging for Alexa-647 showed that kidneys from all embryonic stages had IB4-labeled vasculature, demonstrating that the dye cocktail was distributed to the kidney (Fig. S3D-F). Imaging for Alexa-488 revealed clear apical uptake of dextran in tubules from E15.5 (Fig. 3G,H), while only traces of apical uptake could be seen at E14.5 (Fig. 3I,J), and no apical uptake could be seen at E13.5 (Fig. 3K,L). From this analysis, we propose that flow in the PT initiates at approximately E14. To determine if onset of flow coincides with apical to basal relocation of mitochondria, we compared TOMM20 localization in HNF4A+ PTs of dextran-labeled kidney tissues. As expected, mitochondria in E15.5 PTs with apical dextran uptake were basolaterally located (Fig. 3M,N), while mitochondria in early E15.5 PTs without dextran uptake were apically located (Fig. 3O,P). PTs with traces of apical dextran uptake at E14.5 displayed apical localization of mitochondria (Fig. 3Q,R), as did nascent PTs at E13.5 that did not show any evidence of apical dextran uptake (Fig. 3S,T). Co-staining of mitochondria, microtubules and stress fibers in E13.5, E14.5, and E15.5 kidneys revealed that mitochondria in PTs do not colocalize with filamentous actin at any of these developmental stages, while there is some evidence for colocalization of mitochondria and microtubules once the latter form in the E15.5 PT-diff (Fig. S3G-N). Using aquaporin 1 (AQP1) staining to delineate the lumen surface of PTCs we analyzed Z-stacks to confirm that mitochondria are primarily apical in dextran-PT-early (Fig. S3O), while they are abundant basolaterally in dextran+ PT-diff (Fig. S3P). Cross-sections of dextran-PT-early are relatively rare compared with dextran+ PT-diff in the P0 kidney (Fig. S3Q), and in addition to the difference in abundance, there is also a significant difference in epithelial cell morphology. To ask if the basolateral localization of mitochondria correlates with an increase in cytoplasmic volume between PT-early and PT-diff, we compared the cytoplasm:nucleus volume ratio in PT-early versus PT-diff (Fig. S3R). We found that the basolateral relocation of mitochondria indeed does correlate with increased cytoplasmic volume, suggesting that it is a component of the hypertrophy associated with onset of function in the PTC. Our analysis of developmental stages agrees with our observations from newborn kidneys and confirms that PT-early are HNF4A+/dextran-pre-functional tubules with apical mitochondria and PT-diff are HNF4A+/dextran+ functional tubules with basolateral mitochondria.

### Mitochondria are apically located in PTs of kidneys that lack glomeruli

To ask if flow is necessary for mitochondrial relocation, we characterized subcellular localization of mitochondria in a mouse mutant with nephrons that lack glomeruli. *Six2cre*-mediated compound inactivation of the genes for numb and numb-like (*numbl*) results in formation of kidneys that almost completely lack glomeruli, but that still have extensive differentiation of HNF4A+ PTs at E18.5 (Fig. 4A-D, S4A-D). The kidneys of *Six2cre*;*Numb^loxP/loxP^*;*Numbl^loxp/loxp^*(mutant) are similar in size to *Six2cre*;Numb^+/loxP^;*Numbl^+/loxp^*littermate controls (Fig. S4A-D) but have severely attenuated numbers of glomeruli (Fig. 4E, S4A,B). Glomeruli that do form in mutant kidneys are poorly organized, suggesting limited functionality (Fig. 4C). Numbers of nephron tubules in mutant and control kidneys differ only modestly (Fig. 4F, S4C,D), indicating an abundance of aglomerular nephrons in the mutant (the full characterization of the phenotype will be reported in an independent publication). Since the majority of nephrons in mutant kidneys lack glomeruli, they can be used to ask if lumen flow is necessary for basolateral relocation of glomeruli. In control kidneys, NZ-adjacent HNF4A+ PTs display apical mitochondria (Fig. 4G), while those located deeper in the cortex display mitochondria distributed throughout the cytoplasm (Fig. 4H, S4E,F). In contrast, the majority of PTs in both NZ-adjacent and deeper cortex regions of the mutant kidney display apical mitochondria (Fig. 4I,J, S4G,H), and only solitary PTs show evidence of redistribution from this location (Fig. 4J). A small proportion of nephrons in mutant kidneys do have vascularized glomeruli (Fig. 4C), and the small number of PTs with basolateral mitochondria that can be found in mutants likely reflects onset of flow in these rare nephrons.

**Figure 4:**
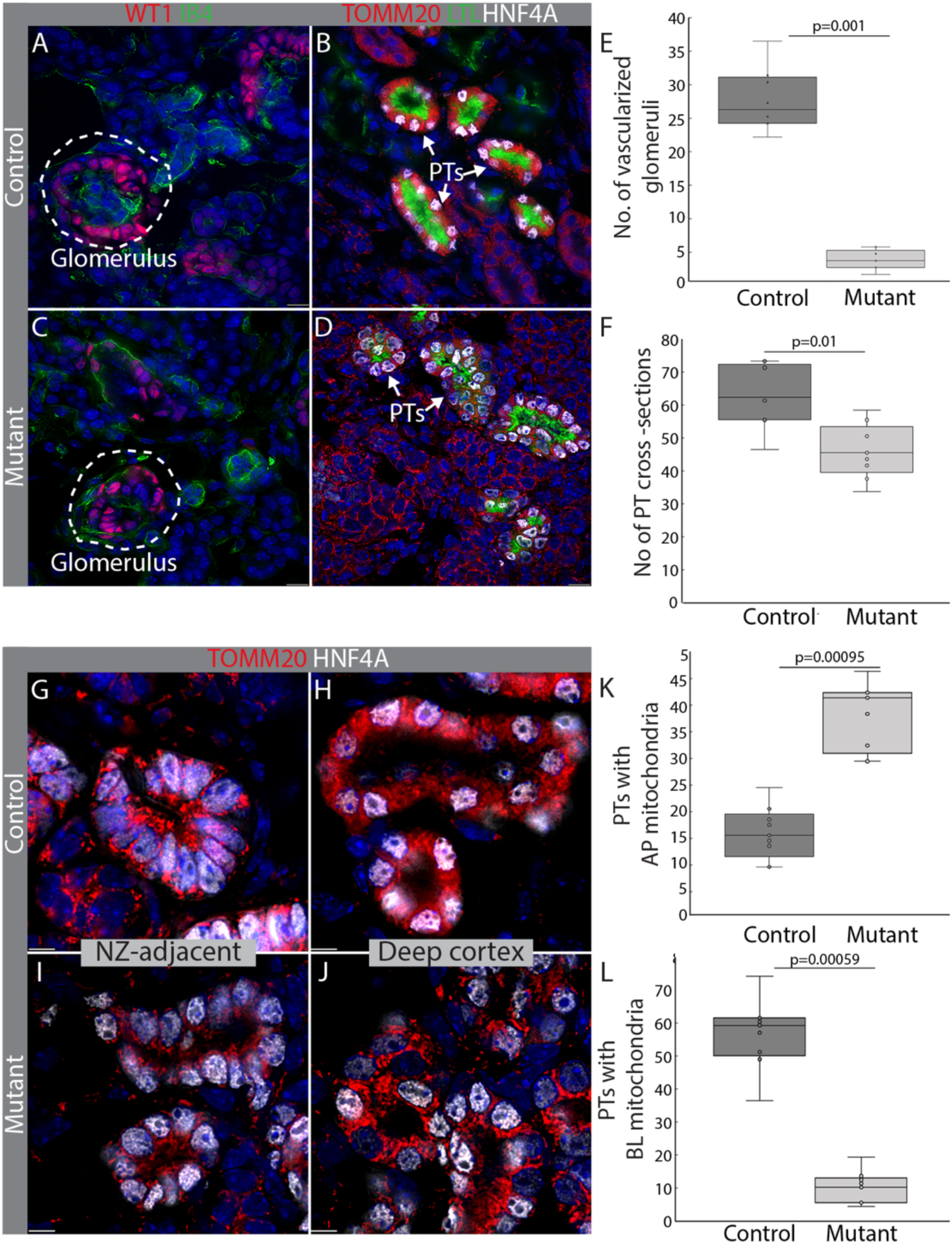
Mitochondria are apically localized in aglomerulur nephrons. E18.5 kidneys from *Six2cre*;Numb^+/loxP^;*Numbl^+/loxp^*(control) and *Six2cre*;*Numb^loxP/loxP^*;*Numb^loxploxp^* (mutant) littermates. (A,C) Identification of glomeruli by staining for WT1 (podocyte marker, red) and IB4 (endothelial cell, green). (B,D) PTs labeled with HNF4A (white) and TOMM20 (mitochondria, red). (E,F) Quantification of glomeruli and PTs in triplicate sections of control kidneys vs triplicate sections of mutant kidneys (N=3 individuals each). Data is shown as box-and-whisker with mean (middle line) and minimum–maximum values (whiskers); two-tailed Student’s *t*-test. (G-J) Control and mutant kidneys stained with HNF4A (PT-white) and TOMM20 (mitochondria-red). Nephrogenic zone-adjacent PTs from control (G) and mutant (I) kidneys showing largely apical mitochondria. PTs from the deep cortex of control kidneys (H) showing largely basolateral mitochondria and from mutant kidneys (J) showing abundant apical mitochondria. (K,L) Quantification of PT localization in sections from control vs mutant (N=3 individuals each) shown as box-and-whisker with mean (middle line) and minimum–maximum values (whiskers); two-tailed Student’s *t*-test. **Abbreviations:** PT, proximal tubule; NZ, nephrogenic zone. All sections were counterstained with the nuclear dye DAPI (blue). Scale bars in (A-D) 10 µm, and (G-J) 5 µm.

Quantification of apical mitochondrial localization is shown in Fig. 4K, and the increase in abundance of apical mitochondria in the mutant correlates well with the reduction in basolateral mitochondria, which is shown in Fig. 4L. This analysis shows that PTs of aglomerular nephrons lacking lumen flow do not redistribute mitochondria from their apical to basolateral aspect. To confirm this observation using an independent PT marker, we co-stained control and mutant kidneys with aquaporin 1 (AQP1), LTL and TOMM20 (Fig. S4I-N’). AQP1 labels membranes of PTCs in NZ-adjacent PT-early weakly in both control (Fig. S4I,J,J’) and mutant (Fig. S4L,M,M’). PT-diff in deeper layers of the cortex in the control show strong AQP1 staining and abundant basolateral mitochondria (Fig. S4I,K,K’), whereas PTs in deeper layers of the mutant cortex exclusively resemble PT-early, maintain weak expression of AQP1 and apical mitochondria (Fig. S4L,N.N’).

### Redistribution of apical mitochondria associates with differentiation of physiological features and reduced proliferation

Fluid flow has been proposed to initiate a program of cellular changes required for epithelial function, including cytoskeleton organization, formation of microvilli, expression of Na-K ATPase and reduced proliferation, and we were interested to determine if redistribution of mitochondria from their apical location associated with this transition. PT-early tubules were compared with PT-diff tubules in P0 kidneys from mice that had been injected with 10 kD dextran-Alexa488. While ɑ-tubulin could be detected diffusely throughout the cytoplasm of PT-early cells (Fig. 5A), clear networks of ɑ-tubulin microtubules could be seen in PT-diff cells (Fig. 5B). Quantification of microtubule filament junctions shows a significant difference between PT-early and PT-diff (Fig. 5C). To ask if the lack of microtubule organization in PT could be due to a lack of microtubule organizing center, we stained for γ-tubulin, but found similar signal in PT-early and PT-diff (Fig. S5A-D). Filamentous actin staining with phalloidin revealed weak expression on the lumen aspect of PT-early cells (Fig. 5D), and strong expression in PT-diff cells (Fig. 5E) which correlated well with increased glycocalyx staining by LTL (Fig. S5E-H). Quantification supported a significant increase in staining of actin between PT-early and PT-diff (Fig, 5F). Likewise, basolateral expression of the Na-K ATPase was barely detectable in PT-early cells (Fig. 5G), but clearly detectable in PT-diff cells (Fig. 5H), a difference that was also supported by quantification (Fig. 5I). To understand if these changes are associated with reduced cell proliferation, we pulsed P0 pups 2 hours before harvest with the uridine analog EdU, which can be fluorescently detected in tissue sections (Fig. S5H). While EdU-labeled cells are detected in the majority of sections of PT-early tubules (Fig. 5J), they are rare in PT-diff tubules (Fig. 5K). The drastic reduction in proliferation between PT-early and PT-diff is supported by quantification (Fig. 5L). Overall, this analysis indicates that the apical to basolateral relocation of mitochondria seen from PT-early to PT-diff occurs concomitantly with differentiation of features of epithelial function and deceleration of cellular proliferation, suggesting that mitochondrial relocation is part of a program of developmental hypertrophy triggered by onset of lumen flow. Mitochondria associate with the microtubule network to establish many aspects of their architecture (Fenton et al., 2021), so we were particularly interested in visualizing the colocalization of TOMM20 and α-tubulin in the PT-early to PT-diff transition. In PT-early, both mitochondrial and diffuse microtubule staining can be seen largely apically (Fig. 5M-M’’) while in PT-diff α-tubulin forms a microtubule network extending basolaterally with mitochondria distributed throughout (Fig. 5N-N’’). Our data suggest a model for epithelial differentiation in which lumen flow initiates a program of cellular changes including organization of the microtubule network and concomitant redistribution of apically located mitochondria throughout the cytoplasm (Fig. 5O). Based on our TMRM analysis, this would associate with a strong increase in mitochondrial membrane potential and increased energy production required for physiological function. To develop a system with which we could define a mechanistic basis for apical retention of mitochondria in PT-early, we used organoids, which do not experience lumen flow.

**Figure 5:**
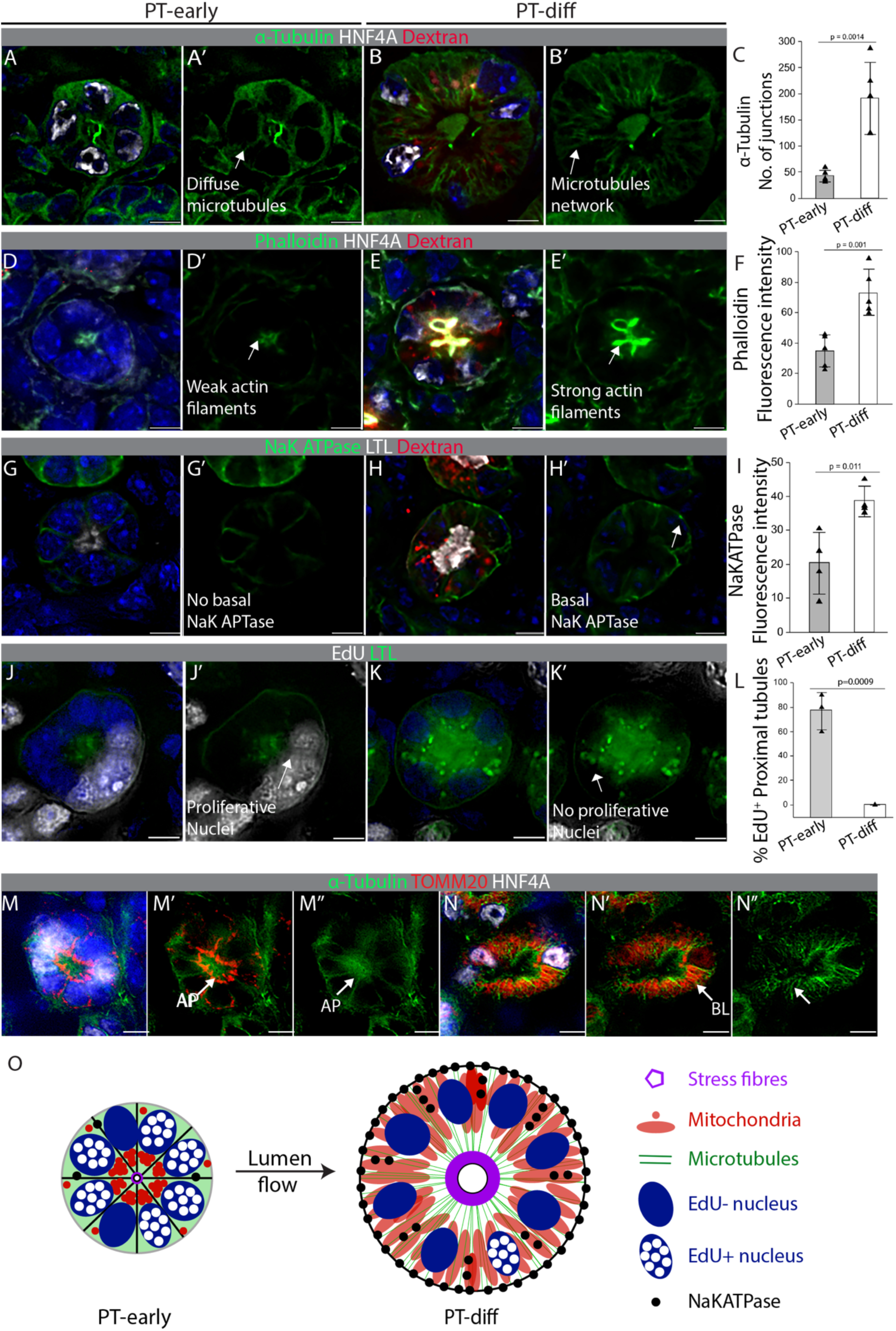
Basolateral localization of mitochondria coincides with microtubule network establishment in PTs. Kidney from P0 mouse pulsed with 10kD dextran-Alexa568 to distinguish between pre-functional, dextran negative PT-early vs functional dextran positive PT-diff. (A-B’) α-tubulin (green) and HNF4A (white). (C) Comparison of number of junctions of α-tubulin in PT-early vs PT-diff. (D-E’) Phalloidin (green) and LTL (white) (n=4). (F) Comparison of abundance of stress fibers in PT-early vs PT-diff. (G-H’) NaK ATPase (green) and LTL (white) (n=4). (I) Comparison of abundance of NaK ATPase pump in PT-early vs PT-diff. (J-K’) EdU (white) and LTL (PT). For EdU labeling studies (n=3 kidney), PT-early were classified as tubules with barely discernible LTL staining, while PT-diff were classified as tubules with strong LTL staining. (L) Comparison of proliferation in PT-early vs PT-diff. (M-N”) Co-localization of α-tubulin (green), TOMM20 (red) and HNF4A (white) in dextran-negative (M-M’’) versus dextran-positive (N-N’’) tubules. (O) Schematic model for organelle redistribution in the PT-early to PT-diff transition that we propose is initiated by lumen flow. All bar graphs tested with two-tailed Student’s *t*-test. **Abbreviations:** EdU, 5-ethynyl deoxyuridine; PT, proximal tubule. All sections were counterstained with the nuclear dye DAPI (blue). Scale bars in all panels 5 µm.

### PT mitochondria in organoids are primarily apically located

Kidney organoid tissues from human pluripotent stem cells show extensive differentiation of PTs, and provide important organotypic model systems for the study of PT biology. To determine the subcellular localization of PT mitochondria in organoids, we analyzed TOMM20 expression in HNF4A+ tubules in organoids generated from H9 human embryonic stem cells cultured using the conditions described in (Morizane et al., 2015). These show extensive differentiation of podocytes, PTs and distal tubules at day 16 of culture (Fig. 6A), and these differentiated cell types are maintained for the 30 days of culture conducted in this study (Fig. S6A-D). Analysis of TOMM20 expression in PTs at 16, 23, and 30 days of culture revealed primarily apical mitochondria (Fig. 6B,S6B,D), and can be classified as PT-early by comparison with fetal human and mouse PTs. To understand if apical to basolateral relocation of mitochondria could relate to culture time, we quantified the ratio of apical:basolateral TOMM20+ mitochondria in PTs from organoids cultured for 16, 23, and 30 days and compared with apical:basolateral ratio in PT-early vs PT-diff from P0 mouse kidneys (Fig. 6C). We found that the ratios of apical:basolateral mitochondria at all three stages of organoid culture compare with ratios seen in P0 PT-early, indicating that time in culture up to 30 days does not promote apical to basolateral relocation of mitochondria and supports the conclusion that PTs in kidney organoids reflect mitochondrial localization seen in the earliest stage of fetal HNF4A+/LTL+ PT (Fig. 1E). To validate our findings in organoids generated from a different stem cell line, we analyzed PTs derived from AICS-0078-79 induced pluripotent stem cells (iPSCs) that express a TOMM20-mEGFP fusion protein and an α-tubulin-mTagRFP-T fusion protein (Viana et al., 2023). Vigorous tubule formation was seen in these kidney organoids, and fluorescent protein imaging confirmed the observation of primarily apically located mitochondria in PTs at 16, 23, and 30 days of culture (Fig. 6D,E,F). A direct comparison of mitochondrial staining in HNF4A+ PT-early from mouse P0 kidneys, human fetal kidneys, and human kidney organoids on day 16 of differentiation shows comparable apical location (Fig. 6G-I). To determine the expression of mitochondrial localization genes in organoids versus kidneys, we compared expression of genes in GO term G0051646 “mitochondrion localization” between available single cell datasets for human fetal kidney (Lindström et al., 2018) (Fig. S6E) and adult human kidney (Wu et al., 2018) (Fig. S6F) versus human kidney organoid (Wu et al., 2018) (Fig. S6G). Since the focus of our study is the PT, we excluded genes that are not expressed in: 1) Tubule precursors of human fetal kidney; 2) PT of human adult kidney; 3) PT of human kidney organoid. As predicted based on prior studies showing that kidney organoids correspond most closely with fetal kidney (Little and Combes, 2019), we found a strong overlap in gene expression between fetal kidney and kidney organoids, with a weaker overlap with adult kidney (Fig. S6G,H). This genetic analysis supports the notion that organoids largely share the mitochondrial localization program of the fetal kidney, in which the PT-early to PT-diff transition that we are interested in modeling occurs.

**Figure 6:**
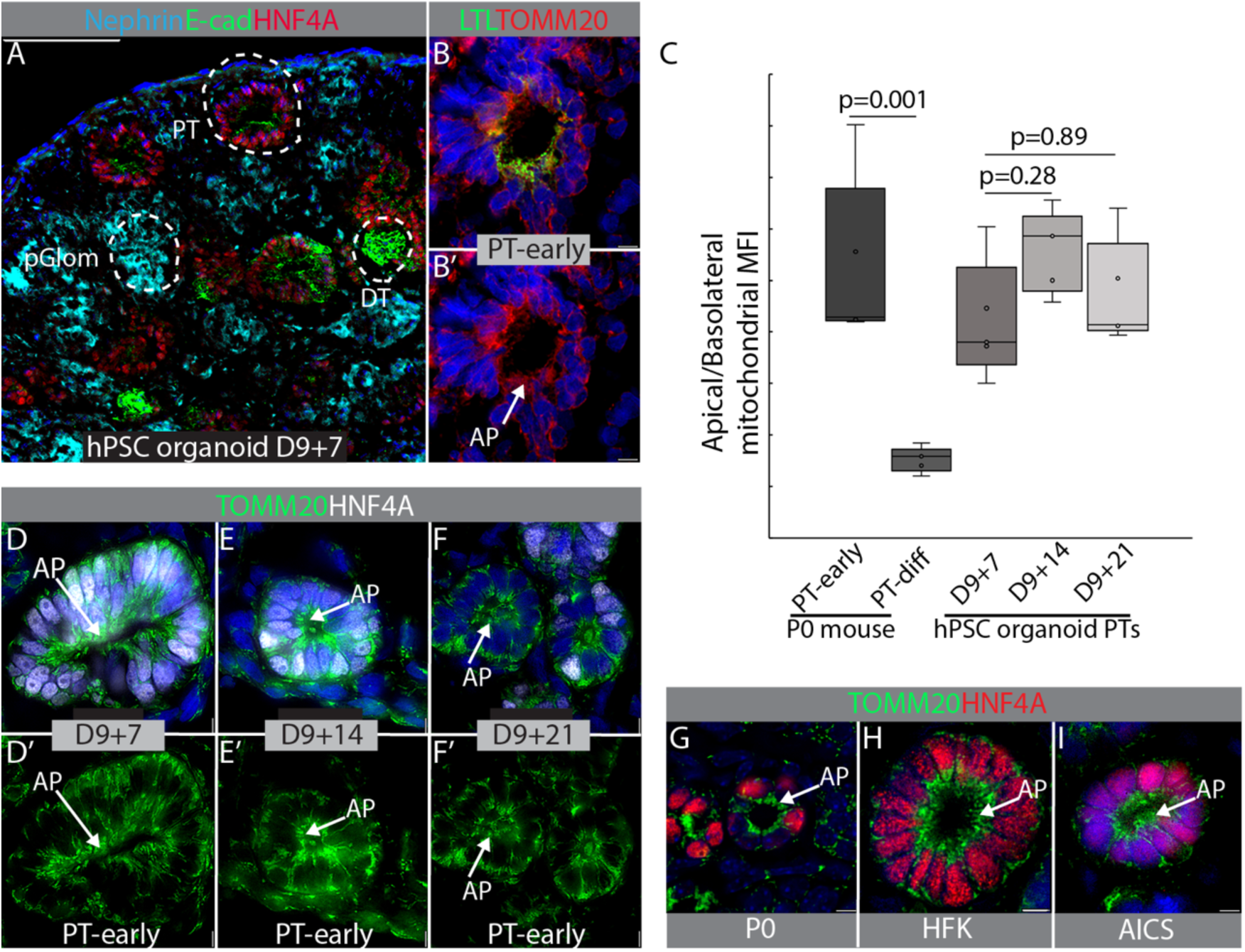
Mitochondria in organoid PTs primarily localize apically. (A) Kidney organoid differentiated from H9 hESCs at 16 days (9+7, 9 days differentiation in adherent cell culture followed by 7 days non-adherent culture of aggregated cells). HNF4A (PT, red); E-Cadherin (distal tubules, green); nephrin (podocytes, cyan). (B) Overlay of PT labeled with LTL (green) and the mitochondrial marker TOMM20 (red), and (B’) TOMM20 alone. (C) Comparison of apical to basolateral TOMM20 in HNF4A+ early and PTs-diff from P0 mouse kidneys (n=5) with HNF4A+ PTs from H9 organoids harvested after 9+7, 9+14 and 9+21 days of culture (n=5 analyzed from each stage). Graph plotted is ratio of apical to basolateral mean fluorescence intensity of TOMM20 in PT. Data is shown as box-and-whisker with mean (middle line) and minimum–maximum values (whiskers); one way ANOVA with post-hoc Tukey HSD test. (D-F) Fluorescent protein detection in organoids from AICS-0078-79 iPSCs that express a TOMM20-GFP fusion protein (green) counterstained with HNF4A (white). (D, D’) Day 9+7, (E, E’) Day 9+14, and (F, F’) Day 9+21. (G-I) TOMM20 (green) and HNF4A (red) in PT-early of (G) P0 mouse PT-early, (H) HFK PT-early and (I) AICS-0078-79-derived kidney organoids on day 9+7 of culture. **Abbreviations:** AP, apical; DT, distal tubule; pGlom, presumptive glomerulus; PT, proximal tubule. All immunofluorescence images are counterstained with the nuclear stain DAPI (blue). Scale bar in A 100 µM, all other panels 5 µM.

### Inhibition of LRRK2 promotes basolateral localization of mitochondria in organoid PTs

Having established that PTs in organoids have mitochondrial localization corresponding to fetal PTs prior to onset of tubule function, we decided to use them as an experimental system to identify molecular pathways that control localization of mitochondria in the prefunctional PT. Review of the literature revealed several factors that regulate subcellular localization of mitochondria in cells including establishment of an actin network (Basu et al., 2021), glutamate signaling (Macaskill et al., 2009), signaling from the primary cilium (Power et al., 2024), and activity of the multifunctional kinase LRRK2 (Hsieh et al., 2016). Organoids with fluorescently labeled mitochondria (AICS-0078-79) were treated with a panel of small molecules targeting these pathways used at their maximal tolerated doses (Fig. S7A-E’), and the apical:basal ratio of mitochondria in PTs was calculated (Fig. 7A, S7M-R). To minimize the risk that secondary effects of pathway modulation would confound our analysis, we limited the treatment period to 4 hours. Inhibitors of two pathways showed an effect on mitochondrial localization: the NMDA receptor and the multifunctional kinase LRRK2. Addition of NMDA to the culture medium had no effect, whereas inhibition of the receptor with dizocilpine (MK-801) showed a significant increase in basolateral localization of mitochondria. This may be explained by the presence of 20 mg/l glutamate in the basal medium (advanced RPMI) which could be expected to cause a basal state of NMDA receptor activation. Analysis of publicly available single cell transcriptomic data revealed LRRK2 is expressed specifically in tubule precursors of human fetal kidney and PTs of adult human kidney (Fig. S6E-G), leading us to focus on this candidate since it may represent a PT-specific pathway for mitochondrial localization. Immunostaining of kidney organoids derived from iPSCs that express YFP under the control of the HNF4A locus revealed LRRK2 in cytoplasm of PTCs (Fig. 7B), but little or no expression in glomeruli or HNF4A-negative presumptive distal tubules (Fig. 7C). Similarly, co-staining of P0 kidneys with LTL and anti-LRRK2 revealed cytoplasmic expression in lectin-stained PTs (Fig. 7D), and both PT-early (Fig. 7E) and PT-diff (Fig. 7F) were strongly positive for LRRK2. To confirm the initial observation from our panel of inhibitors, we compared treatments with two independent small molecule inhibitors of LRRK2 kinase activity, GSK2587215A (Reith et al., 2012) and MLi2 (Sanz Murillo et al., 2023). Immunoblot to confirm inhibition of LRRK2 phosphorylation revealed a reduction or loss of pLRRK2 following treatment with both inhibitors, while total LRRK2 was unaffected (Fig. S7S,T). Both LRRK2 inhibitors caused basolateral localization of mitochondria in organoid PTs (Fig. 7G-J, S7U-W).

**Figure 7:**
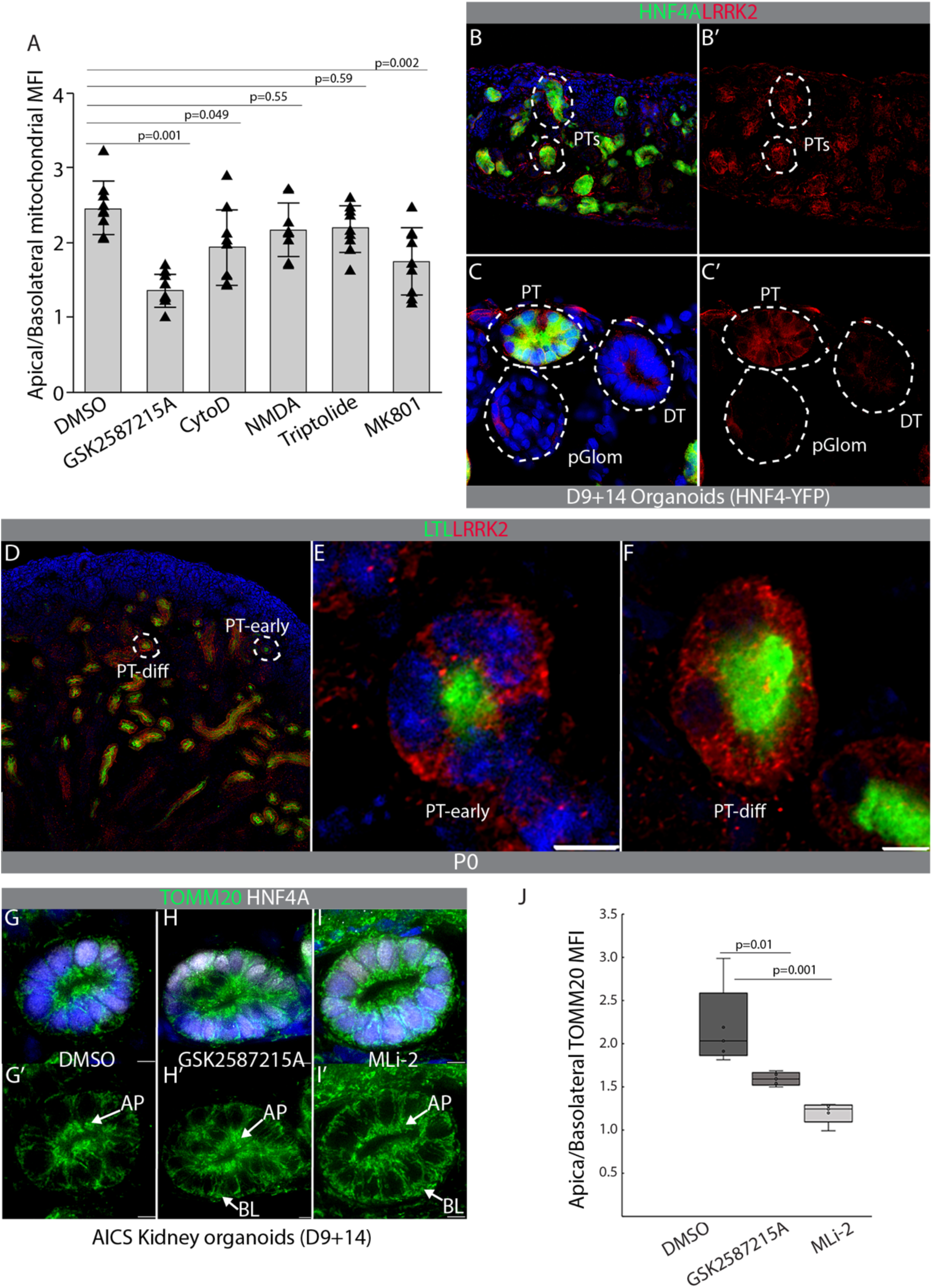
Small molecule panel identifies LRRK2 as a mediator of apical localization of mitochondria in PTs. (A) Ratio of apical to basolateral mitochondria in PTs of kidney organoids derived from AICS-0078-79 iPSCs following 4 hours compound treatment (n=3 organoids), 3 PTs per organoid. (B-C’) D9+14 Kidney organoids derived from HNF4A-YFP iPSCs (YFP under HNF4A promoter, PTs (green)) immunostained for LRRK2 (red). (D-F) P0 mouse kidneys were stained for LTL (PT, green) and LRRK2 (red). (D) Widefield of P0 mouse kidney showing LRRK2 (red) expression restricted to PTs. (E) PT-early displaying cytoplasmic LRRK2. (F) PT-diff displaying cytoplasmic LRRK2. (G-I) Maximum intensity projections of PTs from AICS-0078-79 kidney organoids (D9+14) with eGFP tagged mitochondria (green) stained with HNF4A (PTs, white) following 4 hours treatment with (G-G’) vehicle, (H-H’) GSK2587215A, and (I-I’) MLi-2. (J) Comparison of apical:basolateral ratio of mean fluorescence intensity of TOMM20/PT (n=5 per condition). Data is shown as box-and-whisker with mean (middle line) and minimum–maximum values (whiskers); one way ANOVA with post-hoc Tukey HSD test. **Abbreviations:** AP, apical; BL, basolateral; DT, distal tubule; pGlom, presumptive glomerulus; PT, proximal tubule. All immunofluorescence images are counterstained with the nuclear stain DAPI (blue). Scale bar in D 100 µm, all other panels 5 µm.

### Inhibition of LRRK2 promotes basolateral localization of mitochondria in prefunctional PTs *in vivo*

Based on its selective expression in PTs at various stages of differentiation in kidneys and organoids, and the basolateral localization of mitochondria seen after its inhibition, we hypothesized that LRRK2 controls apical localization of mitochondria in the pre-functional PT. To critically test this hypothesis, we investigated mitochondrial localization following LRRK2 inhibition *in vivo*. Based on multiple reports of availability and selectivity of MLi-2 in rodents (Albanese et al., 2021; Bryce et al., 2021; Bu et al., 2023; Fell et al., 2015; Kluss et al., 2021; Mutti et al., 2023), we selected this inhibitor for *in vivo* studies. P0 mice were intraperitoneally injected 18 hours and 4 hours before harvest with 30 mg/kg MLi-2. Immunoblot for the active phosphorylated form of LRRK2 showed efficient inactivation in kidneys from 5 mice, while total LRRK2 was maintained (Fig. 8A,B). Co-staining for mitochondria and the PT marker HNF4A revealed that LRRK2 inhibition caused a significant loss of apical mitochondrial localization in PT-early (Fig. 8C-E). To ask if LRRK2 inhibition led to energetic consequences for cells that might confound our analysis of mitochondrial localization, we compared incorporation of the mitochondrial dye Mitoview Green and the mitochondrial membrane potential-dependent dye TMRM in hTert-RPTEC treated with vehicle versus MLi-2 (Fig. S8A-F). To confirm that TMRM was reporting on mitochondrial membrane potential and energetic output, we measured TMRM signal after treatment with the mitochondrial membrane depolarizer CCCP (Fig. S8C,F). Quantification revealed no significant difference between vehicle and MLi-2 treatment for Mitoview (Fig. S8G) or TMRM (Fig. S8H), and CCCP treatment showed that TMRM signal is indeed dependent on mitochondrial membrane potential (Fig. S8H). We concluded that a change in energetic state of mitochondria in MLi-2 treated cells is not a significant confounder in our experiments. For genetic confirmation of the observation that LRRK2 inhibition perturbs mitochondrial localization in PT-early, we analyzed the *Lrrk2* loss of function mouse (Tong et al., 2010). *Lrrk2* null kidneys display PT differentiation comparable to wild type littermate controls (Fig. S8I,J). However, analysis of TOMM20 in PT-early of *Lrrk2* null kidneys revealed increased basolateral localization of mitochondria compared with wild type littermate controls (Fig. 8F-H). The concordance between mitochondrial phenotypes observed following pharmacological and genetic inactivation suggests that LRRK2’s kinase activity is required for apical retention of mitochondria in PT-early. Mitochondrial mislocalization also appears to be a feature of PT-diff in *Lrrk2* null P0 kidneys, with regions of PTC cytoplasm lacking TOMM20+ mitochondria (Fig. S8K,L) that resemble vacuoles in shape. PT-diff from MLi-2 treated P0 kidneys display the same changes in mitochondrial localization as the *Lrrk2* mutant (Fig. S8M,N). Reports characterizing this mutant strain and a strain with a kinase dead variant of *Lrrk2* described vacuolation of PT and accelerated aging that initiates in the adult (Herzig et al., 2011; Tong et al., 2010), but our findings suggest that this loss of function phenotype arises already during fetal development.

**Figure 8:**
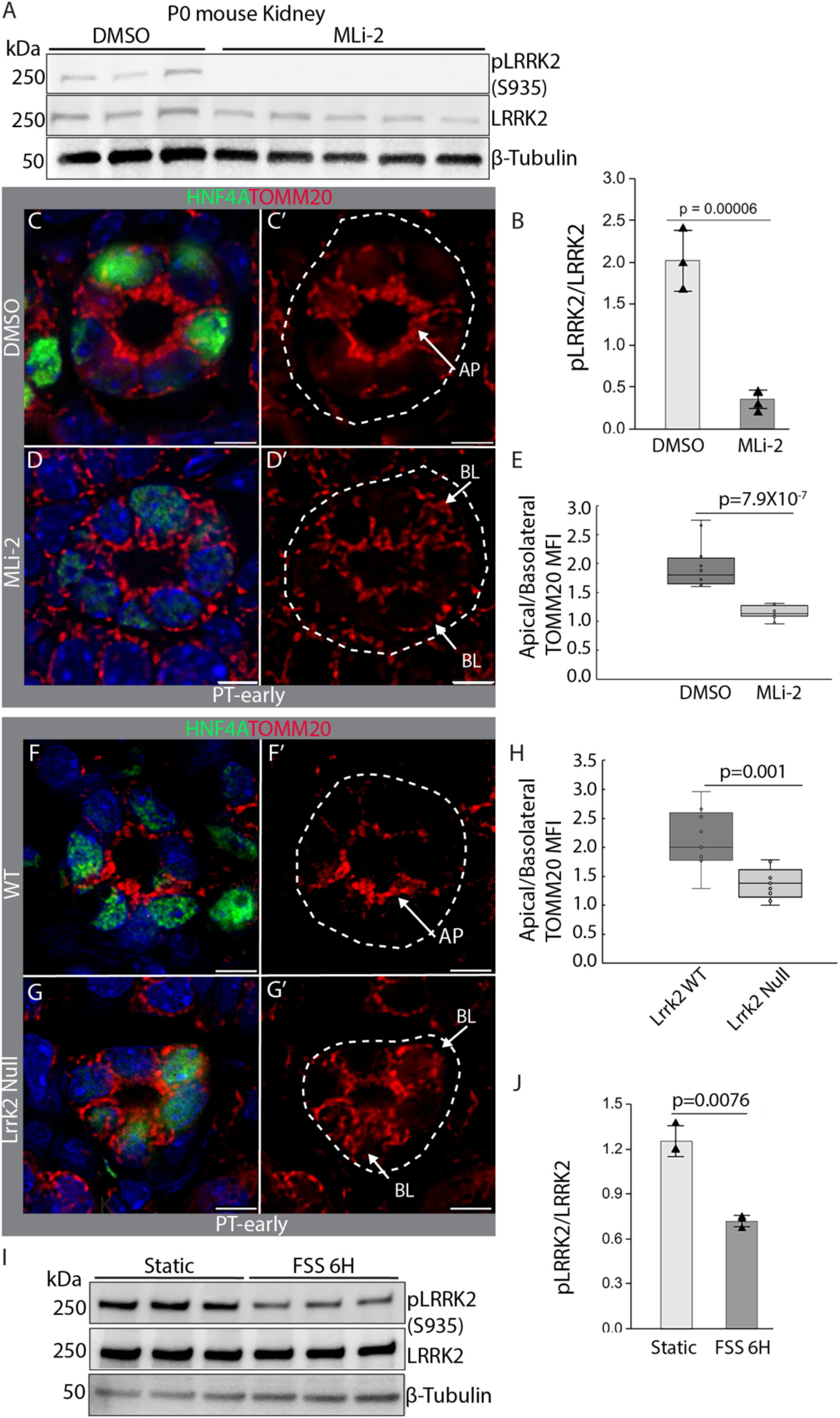
LRRK2 controls apical localization of mitochondria in the newborn kidney. (A) Immunoblot of kidney lysates from P0 mice treated with vehicle (DMSO) or 30 mg/kg MLi-2 (MLi-2) blotted for active phosphorylated LRRK2 (pLRRK2, S935), total LRRK2 and loading control (ꞵ-tubulin). (B) Densitometric quantification of immunoblot plotted as bar graph; two-tailed, unpaired Student’s *t*-test. (C-D’) PT-early from P0 mouse pulsed with 10 kD dextran-A647 stained with HNF4A (PT, green) and TOMM20 (mitochondria, red). (C-C’) DMSO treated mouse. (D-D’) MLi-2 treated mouse. (E) Comparison of apical:basolateral ratio of mitochondria in PT-early from 3 kidneys per condition DMSO-treated (n=4 PT/kidney) versus MLi-2-treated (n=4 PT/kidney). Data is shown as box-and-whisker with mean (middle line) and minimum–maximum values (whiskers); two-tailed, paired Student’s *t*-test. (F-G’) Representative wild type (WT) and *Lrrk2* null P0 mouse littermate kidneys from analysis of 3/group pulsed with 10kD dextran-A647 and stained for HNF4A (PT, green) and TOMM20 (mitochondria, red). (F-F’) PT-early from WT. (G-G’) PT-early from *Lrrk2* null. (H) Comparison of apical:basolateral ratio of mitochondria in PT-early in kidneys from 3 individuals per cohort of WT (n=3PT/kidney) versus *Lrrk2* null (n=3PT/kidney). Data is shown as box-and-whisker with mean (middle line) and minimum– maximum values (whiskers); two-tailed unpaired, Student’s *t*-test. (I) Immunoblot of hTert-RPTEC subjected to fluid shear stress (FSS) for 6 hours and blotted for phosphorylated LRRK2 (pLRRK2-S935), total LRRK2 and loading control (ꞵ-tubulin). (J) Densitometric quantification of immunoblot plotted as bar graph; two-tailed unpaired Student’s *t*-test. **Abbreviations:** AP, apical; BL, basolateral. PT, proximal tubule. Immunofluorescence images C and E are counterstained with the nuclear stain DAPI (blue).

### Activity of LRRK2 is modulated by fluid shear stress

The observation that pharmacological inhibition of LRRK2 in P0 mice relocates mitochondria from their apical position in PT-early suggests that the transition from PT-early to PT-diff is accompanied by a reduction in LRRK2 activity. Immunostaining for LRRK2 shows that the protein is expressed both in PT-early and PT-diff, and thus a change in overall protein expression does not explain the reduction in activity. Analyses of dextran uptake in embryonic kidneys (Fig. 3) and mutant kidneys lacking flow (Fig. 4) support the hypothesis that onset of lumen flow triggers mitochondrial relocation, and based on this we reasoned that flow may control LRRK2 activity. To test this, we subjected confluent monolayers of hTert-RPTEC PTCs to fluid shear stress using an orbital shaker, a technique developed for the study of fluid shear stress in endothelium (Kraiss et al., 2003) that has been adapted for kidney (Long et al., 2017). We found that subjecting PTCs to only 6 hours of fluid shear at a rate of 146 revolutions per minute resulted in significant down-regulation of LRRK2 phosphorylation, while LRRK2 protein abundance remained unchanged (Fig. 8I,J). From this experiment we conclude that fluid shear stress indeed controls the activity of LRRK2, providing an explanation for how the transition from PT-early to PT-diff may associate with reduction of LRRK2 activity and redistribution of mitochondria from their apical position.

## Discussion

Formation of the cellular machinery to drive energy production is an essential step in PTC differentiation, ensuring that the extreme energy demands placed on this cell can be met when function ensues. This report focuses on how mitochondria that are located apically in the nascent and immature PTC are redistributed throughout the cell to reach their ultimate basolateral location (Fig. 1). Analysis of mitochondrial membrane potential *in situ* indicates that apically located mitochondria are relatively depolarized, suggesting that their energy output is modest (Fig. 2), and that the transition from apical to basolateral localization associates with an increase in energy output. By defining pre-functional versus functional PTs through measuring their uptake of dextran finding that is supported by analysis of PTC uptake of dextran, we provide evidence that onset of lumen flow triggers the basolateral relocation of mitochondria (Fig. 3), a model that is supported by analysis of mice that are genetically deficient in glomeruli (Fig. 4). Molecular marker analysis indicates that mitochondrial redistribution is one component of a cellular program in the nascent PTC that includes cessation of proliferation, establishment of a microtubule network, development of an apical actin network, and basolateral localization of the Na-K ATPase that ensures appropriate subcellular localization of organelles and forms a template for the differentiated PTC prior to the explosive increase in biogenesis that is characteristic of this cell type’s maturation (Fig. 5). Using the kidney organoid system, which lacks lumen flow and provides a useful model of the nascent PTC (Fig. 6), we show that mitochondria are actively apically sequestered in these cells, and that this depends on the activity of LRRK2 *in vitro* (Fig. 7) and both pharmacological inhibition and genetic knockout confirm that this is also the case *in vivo* (Fig. 8). Modeling the effect of fluid flow on PTCs demonstrates that LRRK2 activity is downregulated by fluid shear stress (Fig. 8), providing an explanation for how onset of flow in the newly differentiated PT can trigger the apical-to-basolateral dissemination of mitochondria that forms the template for subsequent PTC maturation.

Positioning of mitochondria within cells ensures that energy is produced in physical proximity to energy-demanding processes. This has been most extensively studied in neurons, where the cellular machinery that transports mitochondria to drive energy-demanding systems at subcellular locations remote from the cell body has been characterized (Misgeld and Schwarz, 2017). The mature PTC has a high energy requirement, and demanding systems such as the Na-K ATPase are located in the basal plasma membrane of these cells (Kashgarian et al., 1985). The PTC develops extensive invaginations of the basal plasma membrane (Windhager, 1992), along which mitochondria align in a perpendicular orientation to the basement membrane and form networks (Bergeron et al., 1980; Gaffiero et al., 1983). In this arrangement, mitochondria are wrapped in basal plasma membrane and directly adjacent to Na-K ATPases, ensuring efficient energy transfer. Our findings support a central role for LRRK2 in subcellular positioning of mitochondria, suggesting overlap between the PTC and the neuron in use of pathways that control organelle positioning. Because of the significance of activating mutations of LRRK2 as a cause of Parkinson’s disease (Healy et al., 2008), the majority of reports focus on effects of hyperactive LRRK2 in neurons, and there are relatively few loss of function studies. Given this caveat, LRRK2 has been identified as a central organizer of organelle localization that acts through multiple distinct cellular mechanisms. Two LRRK2-driven organelle transport functions are of particular relevance to the subcellular localization of mitochondria, and may be candidates for the control of mitochondrial localization that we report in this study. Hyper-activated LRRK2 has been shown to bind directly to microtubules and prevent dynein and kinesin-driven movement of organelles along these subcellular tracks (Deniston et al., 2020; Kett et al., 2012). However the conformational change in LRRK2 that causes its close association with microtubules is promoted by the inhibitor MLi-2 (Deniston et al., 2020), and considering that we see an increase in apical to basolateral movement of mitochondria upon MLi-2 treatment, this seems an unlikely explanation. LRRK2s effects on the association of mitochondria with the microtubule network may be a more plausible explanation for the effect described in this study. The small GTPases Miro1/2 (Rhot1/2) physically associate with the outer leaflet of the mitochondrion and form a complex with TRAK1/2 which connects the organelle to the microtubule network and kinesins and dyneins which propel them to their destination (Fenton et al., 2021). LRRK2 can physically interact in this complex and disrupt the association between the organelles and the microtubule network, thus preventing mitochondrial movement (Hsieh et al., 2016). Transcriptional atlases of the PTC and of kidney organoids indicate that both Rhot1/2 and Trak1/2 are expressed, suggesting that this pathway is plausible. Future loss of function studies in organoids will reveal if the LRRK2-Miro-Trak pathway indeed is required for appropriate localization of mitochondria in the PTC.

Why it would be necessary to have a cellular system to prevent mitochondrial redistribution within the nascent PTC remains a topic of speculation, but the drastic reduction of proliferation in PTCs that is associated with the flow-induced program might suggest that the rapid proliferation of these cells required to generate a sufficient number for morphogenesis of a PT requires cellular systems to ensure maintenance of organelles in a fragmented state that is not membrane associated so that they can be evenly distributed between daughter cells during cell division. Interestingly, there appear to be two phases of proliferation in the differentiating PT. The first is the period between renal vesicle formation and onset of flow which is described in this study, and the second is in the somatic growth phase (Vogetseder et al., 2008). Whether LRRK2 has a role in organelle localization in this second growth phase remains to be determined.

While the majority of studies of LRRK2’s function have focused on explaining how heritable hyper-activation of the protein causes cellular malfunction, development of therapeutic candidates to inhibit the protein’s function have stimulated research on consequences of loss of function. Cellular malfunction in the PTC is a hallmark of *Lrrk2* null mice, with accumulation of lysosomes and degenerative changes starting 6 weeks after birth (Herzig et al., 2011), although we report evidence that cellular changes ensue in the *Lrrk2* null PTC already in the fetal period (Fig. 8,S8). Specific molecular pathways underlying these cellular changes have not been defined, but substrates of LRRK2 have been identified that play important roles in nephron epithelial cells (Steger et al., 2017, 2016). RAB35 is required for primary cilium growth and formation of both tight- and adherens junctions in kidney epithelia (Clearman 2024). RAB29 (RAB7L1) is also required for primary cilium growth, and for vesicle trafficking and function of the Golgi network. *Rab29* knockout phenocopies the loss of *Lrrk2*, and compound mutants largely resemble mutants in either gene, strongly suggesting that they operate in a common pathway (Kuwahara et al., 2016).

How phosphorylation of LRRK2 may be regulated by fluid shear stress is an open question. Mechanisms that regulate LRRK2 activity are poorly understood, but one important factor that has been identified is subcellular localization. The majority of LRRK2 molecules are diffusely distributed as monomers in the cytoplasm, but it is the oligomerized membrane-associated fraction that has the highest kinase activity (Berger et al., 2010). Fluid shear stress has been shown to alter fluidity of cell membranes (Haidekker et al., 2000), but candidate mechanisms linking such changes to membrane association of LRRK2 remain to be described.

In this report we identify a novel function for LRRK2 in positioning mitochondria in newly differentiated proximal tubules at pre- and early postnatal stages. Whether this is a truly distinct mechanism from control of the endolysosomal system which has been implicated in adult-onset PTC dysfunction (Kluss et al., 2021), or if mitochondrial and endolysosomal control may represent two facets of a single organelle localization regulator function of LRRK2 will be the subject of our future inquiries.

## Materials and methods

### Human Tissue

Human fetal kidneys were obtained from the University of Pittsburgh Health Sciences Tissue Bank through an honest broker system after approval from the University of Pittsburgh Institutional Review Board (IRB number: 0702050) and in accordance with the University of Pittsburgh anatomical tissue procurement guidelines.

### Mice

Swiss Webster (Taconic Biosciences, Germantown, NY, USA) were used for molecular marker analysis of wild type mice and for dye incorporation studies. *Numb^tm1Zilj^*, *Numbl^tm1Zili^* (Wilson et al., 2007) and *Six2-TGC^tg/+^*(Kobayashi et al., 2008) were bred to generate *Six2cre*;*Numb^loxP/loxP^*;*Numbl^loxP/loxP^*mutants and *Six2cre*;Numb^+/loxP^;*Numbl^loxP-^* controls. *Lrrk2* knockout mice have a deletion of the promoter region and exon 1 (Tong et al., 2010) and are on a hybrid background (B6;129-*Lrrk2^tm2.1Shn^*/J from The Jackson Laboratory). Animal experiments were reviewed and approved by the Institutional Animal Care and Use Committees (IACUC) of New York Blood Center, where the Rogosin Institute research laboratory is located.

### Cryosectioning and Immunostaining

Organoids and kidney tissue were fixed in 4 % paraformaldehyde (Sigma, St. Louis, MA, USA) for 30 min at room temperature and cryoprotected with a series of sucrose (Sigma, St. Louis, MA, USA), 10 % for 10 minutes, 20 % for 45 minutes or until the tissue sank. Tissue /organoids were then embedded in Tissue -Tec® O.C.T (Thermo Fisher Scientific, Waltham, MA, USA). and cryosectioned. Sections were then fixed briefly for 2 minutes with 4 % PFA and permeabilized with 0.1 % Triton X100 (Sigma, St. Louis, MA, USA) for 10 minutes. Blocking buffer consisting of 5 % donkey or goat serum (Jackson Immunochemicals, West Grove, PA, USA) was added for 30 minutes and then primary antibodies (Table 1) diluted (1:100) in PBST were added for 1 hour. Sections were washed for 30 minutes with PBST and incubated with secondary antibodies (Table 1) diluted (1:200) in PBST for 30 minutes. Unbound / non-specific secondary antibody was washed out for 30 minutes with PBST (Sigma, St. Louis, MA, USA). Immunostained sections were then mounted with VECTASHIELD vibrance antifade mounting medium (Vector Laboratories, Newark, CA, USA). Fluorescent images were acquired as Z-stack using Leica THUNDER Imager. Z-stack images were stacked by 3D visualization tool (LASX generate 3D video) to generate 3D video. For quantification, integrated density of fluorescent images was measured using Image J (National Institutes of Health, Bethesda, Maryland, USA). For each antibody stain, at least three biological and technical replicates were measured, and the mean was plotted with standard deviation (mean ± s.d.).

### Electron microscopy

Samples were fixed over night in 2.5 % glutaraldehyde (Hatfield, PA, USA)/2.0 % paraformaldehyde (Hatfield, PA, USA) buffered with 0.1 M sodium cacodylate (Hatfield, PA, USA)Samples were cut along the longitudinal and transverse planes to generate 4 pieces of tissue. Samples were washed with buffer and then post-fixed in a 2 % osmium tetroxide (Hatfield, PA, USA) solution for 2 hours. Following post-fixation, samples were again washed in buffer and then dehydrated in ethanol (30-100 %) (Hatfield, PA, USA); samples were stained en bloc with uranyl acetate during dehydration in 70 % ethanol. Dehydrated samples were then desiccated with propylene oxide (Hatfield, PA, USA) and initial infiltration was done with a 1:1 propylene oxide:epoxy resin solution (Hatfield, PA, USA). Samples were removed from the 1:1 mixture and infiltrated in pure epoxy resin over night. Following infiltration, samples were embedded in pure epoxy resin in size 00 BEEM capsules and allowed to polymerize for 48 hours at 60 °C. Polymerized blocks were then trimmed with a degreased razor blade, and ultrathin sections were generated using a Diatome diamond knife (Quakertown, PA,USA) and RMC Boeckler Powertome (Tucson, Arizona, USA). Sections were collected on 100-mesh formvar/carbon coated copper grids and contrasted with Uranyless and lead citrate. Sections were imaged on a Tecnai Spirit G2 Biotwin (Hillsboro, Oregon, USA) equipped with an AMT camera and imaging software. Micrograph brightness and contrast were corrected using ImageJ2 version 2.17.0/1.54p.

### Mitochondria quantification

For quantification of mitochondrial abundance, the region of interest was thresholded using ImageJ2 version 2.17.0/1.54p after converting the image into an 8 bit image and integrated density of TOMM20 was measured for each PT. Changes in mitochondrial localization were quantified as ratio of apical to basolateral mean fluorescence intensities of TOMM20. Mitochondria to the north (towards the lumen) of the nucleus were segmented out as apical and the rest were classified as basolateral. Using the free hand selection tool, selections for apical and basolateral regions were created and these files were saved with an extension (.roi). These roi files were then re-opened on the original image to measure mean fluorescence intensities of apical versus basolateral mitochondria separately. The free hand selection tool enabled mitochondrial quantification in tubules with irregular shapes, for example those in organoids. Mitochondrial length in electron micrographs was measured by drawing a line perpendicular to the cristae equivalent to the mitochondrial length with ImageJ2.

### Microtubule quantification

Microtubule complexity was quantified using ImageJ. The microtubules were skeletonized after thresholding. Analyze 2D/3D (skeleton) plugin was then used to count the number of junctions of the microtubules. Number of junctions per tubule obtained from the ImageJ analysis were representing the microtubule complexity in PT-early and PT diff.

### Cytoplasm to Nuclear ratio quantification

Cytoplasm to nuclear ratio was quantified by ImageJ2. Images were stacked and threshold on DAPI (nucleus) AQP1 (cell membrane) was adjusted manually for each condition. 3D object counter plugin was used to calculate cytoplasm and nuclear volume. Data obtained ImageJ is represented as cytoplasm to nuclear ratio in μm^3^.

### TMRM labeling

P0 mice were injected intravenously with 3 mg/kg tetramethylrhodamine methyl ester (TMRM) (ThermoFisher, Waltham, MA, USA) suspended in saline. 20 minutes after injection, kidneys were harvested and imaged within 15 minutes. Alternatively, TMRM loading was performed by immersion of bisected kidneys within 2 minutes of euthanasia in a solution of Advanced DMEM containing 50 µg/ml TMRM for 7 minutes at room temperature on a Nutator. Stained tissues were rinsed briefly in PBS, mounted onto glass coverslips, and imaged on Leica Thunder Imager. To study the influence of efflux pumps on TMRM fluorescence, 2 mM of the p-glycoprotein inhibitor verapamil (ThermoFisher, Waltham, MA, USA) (Wang and Sun, 2020) was added to the staining solution. To gauge the level of tissue autofluorescence in the assay, 1 mM of the mitochondrial membrane depolarizer CCCP (Sigma, St. Louis, MA, USA) was added to the staining solution. The background signal of CCCP-treated sections was subtracted to generate a heatmap of TMRM fluorescence as a proxy measurement of mitochondrial membrane polarization in the P0 mouse kidney.

### Dextran labeling

FITC or Alexafluor-647 labeled 10 kD dextran (8 mg/kg) (Thermo Fisher Scientific, Waltham, MA, USA) was injected retro-orbitally in mouse embryos and newborns. For injection of newborn mice, retro-orbital injection of 25 µl dextran solution was performed rapidly and the pups were returned to the dam. For injection of embryos, uteri were dissected out of pregnant females and the fetal membranes containing the embryos were ruptured without severing the umbilicus. Retro-orbital injection of 5 µl dextran solution was then performed and vascular distribution of dextran was confirmed after one minute by fluorescence microscopy of the hindlimbs. Embryos that did not display intravascular fluorescent dextran were severed from the uterus. The uterus with successfully injected embryos attached to their placentae by the umbilical cord was then incubated in 37 °C Advanced DMEM (ThermoFisher, Waltham, MA, USA). As a tracer to validate dye delivery to embryonic kidneys, Alexafluor-647-labeled Griffonia simplificata isolectin B4 (IB4) (Thermo Fisher Scientific, Waltham, MA, USA) was included in the dextran solution at 1 mg/kg. Kidneys were harvested 30 minutes after injection, fixed, cryosectioned, immunostained and imaged.

### EdU labeling

P0 mice were intraperitoneally injected with 200 mg/kg of 5-ethynyl-2’-deoxyuridine (EdU, Thermo Fisher Scientific, Waltham, MA, USA). Kidneys were harvested, fixed, and cryosectioned 2 hours after injection with EdU. EdU retaining cells were detected by EdU click-it kit (Thermo Fisher Scientific, Waltham, MA, USA).

### Cell culture and directed differentiation

hTERT-RPTECs (ATCC, Virginia, Washington DC, USA) cells were maintained in hTERT Immortalized RPTEC Growth Kit (ATCC, Virginia, Washington DC, USA). Human iPSC line AICS-0078-79 (Viana et al., 2023) was maintained in mTeSR plus on Matrigel (Corning, Corning, NY, USA), HNF4A-YFP iPSCs (Vanslambrouck et al., 2019) were maintained in StemFit® Basic04 on Matrigel (Corning, Corning, NY, USA), Human H9-SIX2-GFP (this study) were maintained in StemFit® Basic04 (Ajinomoto, Itasca, IL, USA) on Geltrex (Thermo Fisher Scientific, Waltham, MA, USA). All iPSC lines were passaged every 3-4 day with TrypLE Express (Thermo Fisher Scientific, Waltham, MA, USA) as single cells with 10 μM Rho kinase inhibitor, Y27632 (EMD Millipore, Burlington, MA, USA).

hPSCs/ES lines were differentiated to kidney progenitors according to (Morizane et al., 2015). For directed differentiation AICS-0078-79 were plated at a density of 1.02 × 10 ^3^ cells/cm^2^, H9-SIX2 - GFP cells were plated at a density of 9.4 × 10^3^ cells/cm^2^ in a six-well plate and HNF4-YFP iPS cells were plated at a density of 1.02 × 10 ^3^ cells/cm^2^. On day 3, at approximately 40-50 % confluency, medium was replaced with Advanced RPMI 1640 (Thermo Fisher Scientific, Waltham, MA, USA), 1× GlutaMAX^TM^ (Thermo Fisher Scientific, Waltham, MA, USA),with 10 μM CHIR99021 (Reprocell, Beltsville, MD, USA), and 7.5 ng/ml Noggin (PeproTech, Cranbury, NJ, USA) for AICS-0078-79, 9 μM CHIR99021for H9-SIX2 -GFP, and 8 μM CHIR99021 for HNF4-YFP for next 4 days to achieve late primitive streak stage. On Day 4, medium was replaced with advanced RPMI 1640 containing 10 ng/ml Activin A (R&D, Minneapolis, MN, USA) and 1X GlutaMAX^TM^ for next 3 days to differentiate them to posterior intermediate mesoderm. On day 7, medium was replaced with advanced RPMI 1640 containing 10 ng/ml FGF9 (R&D, Minneapolis, MN, USA) and 1X GlutaMAX^TM^ for the next 2 days. On day 9 of directed differentiation, differentiated cells were designated “kidney progenitor cells or metanephric mesenchyme”. Kidney progenitor cells were dissociated with TrypLE Express and resuspended at a density of 1 × 10^5^ cells/μl in Advanced RPMI 1640, 1× GlutaMAX^TM^ medium supplemented with 10 ng/ml FGF9. For making liquid air interface kidney organoids, isopore membranes (EMD Millipore, Burlington, MA, USA) were floated on 1 ml of FGF9 supplemented Advanced RPMI 1640, 1× GlutaMAX^TM^ medium. Resuspended cells were spotted as organoids on top of the floating membranes (2 μl/aggregate). 48 hours after organoids spotting, medium was refreshed. thereafter organoids were maintained in advanced RPMI 1640, 1× GlutaMAX^TM^. For submerged kidney organoids, kidney progenitor cells were resuspended at a density of 5 x 10^5^ cells/ml in Advanced RPMI 1640, 1× GlutaMAX^TM^ medium supplemented with 10 ng/ml FGF9 medium. 200 μl of cell suspension was seeded in each well of ultra-low attachment 96 well plate and centrifuged for 15 seconds at 300 g to allow them to aggregate as small organoids. 48 hours after aggregating organoids, medium was refreshed. After day 4 advanced RPMI 1640, 1× GlutaMAX^TM^ was used as organoids maintenance medium.

### Organoid and cell treatments

For compound treatment of organoids, medium was supplemented with Cytochalasin D (1 µM), GSK2587215A (10 µM), MK801 (1 µM), NMDA C1 (10 µM), Triptolide C1 (0.05 µM) for 4 hours. For LRRK2 inhibition experiments, organoid medium was supplemented with 10 µM of GSK2587215A and 10 µM of MLi-2 and RPTEC medium was supplemented with 5 µM of GSK2587215A and 5 µM of MLi-2 for 4 hours. Small molecule concentrations were the maximal tolerated dose based on changes in organoid morphology (Fig. S7A-E’’) and cell toxicity assays (Fig. S7G-L’). In cell toxicity assays, organoids pretreated with small molecules and stained live with DAPI (1 μg/ml) for 10 minutes 37 °C to visualize dead cells (Fig. S7G-L’). Small molecules were purchased from Tocris Bioscience (Minneapolis, MN, USA).

### Immunoblotting

Cells were lysed with RIPA buffer (Thermo Fisher Scientific, Waltham, MA, USA), 1X cOmplete mini EDTA-free protease inhibitor cocktail (Roche, Indianapolis, IN, USA) and 1x phosphoSTOP (Roche, Indianapolis, IN, USA) as per manufacturer’s instructions. Lysed cells were centrifuged at 12,000 rpm for 30 min at 4 °C. The supernatant was collected as whole cell lysate. Protein concentration was measured by BCA assay (Thermo Fisher Scientific, Waltham, MA, USA), according to manufacturer’s instructions. 15-20 µg of total protein was resolved on 4-15 % mini-PROTEAN TGX precast protein gel (Bio-Rad, Hercules, CA, USA) at 200 V. Proteins were transferred to the nitrocellulose membrane at 1.3 A constant current, for 15 min. After protein transfer, the membranes were blocked with 5 % non-fat dry milk (Sigma, St. Louis, MA, USA) for 1 hour. Membranes were probed with primary antibody (1:2000) diluted in 5 % non-fat dry milk (1X TBST) for 1.5 hours at room temperature. Non-specific primary antibody binding was washed off with 1X TBST, three times for 10 minutes each. Membranes were then incubated with HRP conjugated secondary antibody (1:10000) and strepTactin HRP conjugated (Bio-Rad, Hercules, CA, USA) (1:10000) diluted in 5 % non-fat dry milk (1X TBST) for 1 hour followed by 3 washes of 1X TBST for 10 minutes each. The blots were developed using ECL kit (Thermo Fisher Scientific, Waltham, MA, USA).

### Fluid Shear stress experiment

12 mm coverslips were immobilized with 1 ml of 5% low melt agarose in each well of a 12-well dish and allowed to solidify for 30 minutes. Immobile coverslips were then coated with a 1:100 dilution of Matrigel (Corning, Corning, NY, USA) in cell culture medium for 1 hour. The Matrigel solution was aspirated and hTERT-RPTECs were seeded onto the coverslips at a density of 2.5 x 10^5^ per well with 1 ml of medium (Wieser et al., 2008). Medium was replaced 16 hours after seeding the cells to remove floating cells and cells were allowed to grow further for 72 hours to reach confluence. Thereafter the medium was changed every 48 hours. On reaching confluence, plates were then transferred to an orbital shaker in the incubator and rotated at a speed of 146 rpm (based on (Long et al., 2017)) for 6 hours.

## Abbreviations

NZ: nephrogenic zone;
PT: proximal tubule;
PTC: proximal tubule epithelial cell

## Acknowledgments

Many thanks to the Electron and Confocal Microscopy core facility, LFKRI, New York Blood Center, New York, NY. Work performed in the Pitt Biospecimen Core (RRID:SCR_025229) and services and instruments used in this project were supported, in part, by the University of Pittsburgh, the Office of the Senior Vice Chancellor for Health Sciences.

## Author contributions

Conceptualization: L.O.; Methodology: M.A.K, K.B, L.O.; Validation: L.O.; Formal analysis: M.A.K., L.O.; Investigation: M.A.K., K.B., E.G., D.C, L.Y., L.O.; Resources: L.O., T.C, A.P.M., S.S.L.; Data curation: M.A.K., L.O.; Writing - original draft: M.A.K., L.O.; Writing - review & editing: L.O.; Visualization: M.A.K., L.O.; Supervision: L.O.; Project administration: L.O.; Funding acquisition: L.O., T.C.

