## Supplemental figures 1-8 for "Mitochondrial organization in the developing proximal tubule is controlled by LRRK2"

[1] The Rogosin Institute, 310 E 67th Street, New York, NY 10065, USA
[2] Graduate School of Biomedical Sciences and Engineering, University of Maine, 5775 Stodder Hall, Orono, ME 04469, USA.
[3] Department of Molecular Biology and Internal Medicine, University of Texas Southwestern Medical Center, Dallas, TX 75390, USA.
[4] Department of Pediatrics, University of Pittsburgh Medical Center Children's Hospital of Pittsburgh (UPMC CHP), University of Pittsburgh, Pittsburgh, PA, USA
[5] Department of Stem Cell Biology and Regenerative Medicine, Eli and Edythe Broad Center for Regenerative Medicine and Stem Cell Research, Keck School of Medicine, University of Southern California, Los Angeles, CA 90033, USA. Current address: Division of Biology and Biological Engineering, California Institute of Technology, Pasadena, CA 91125.

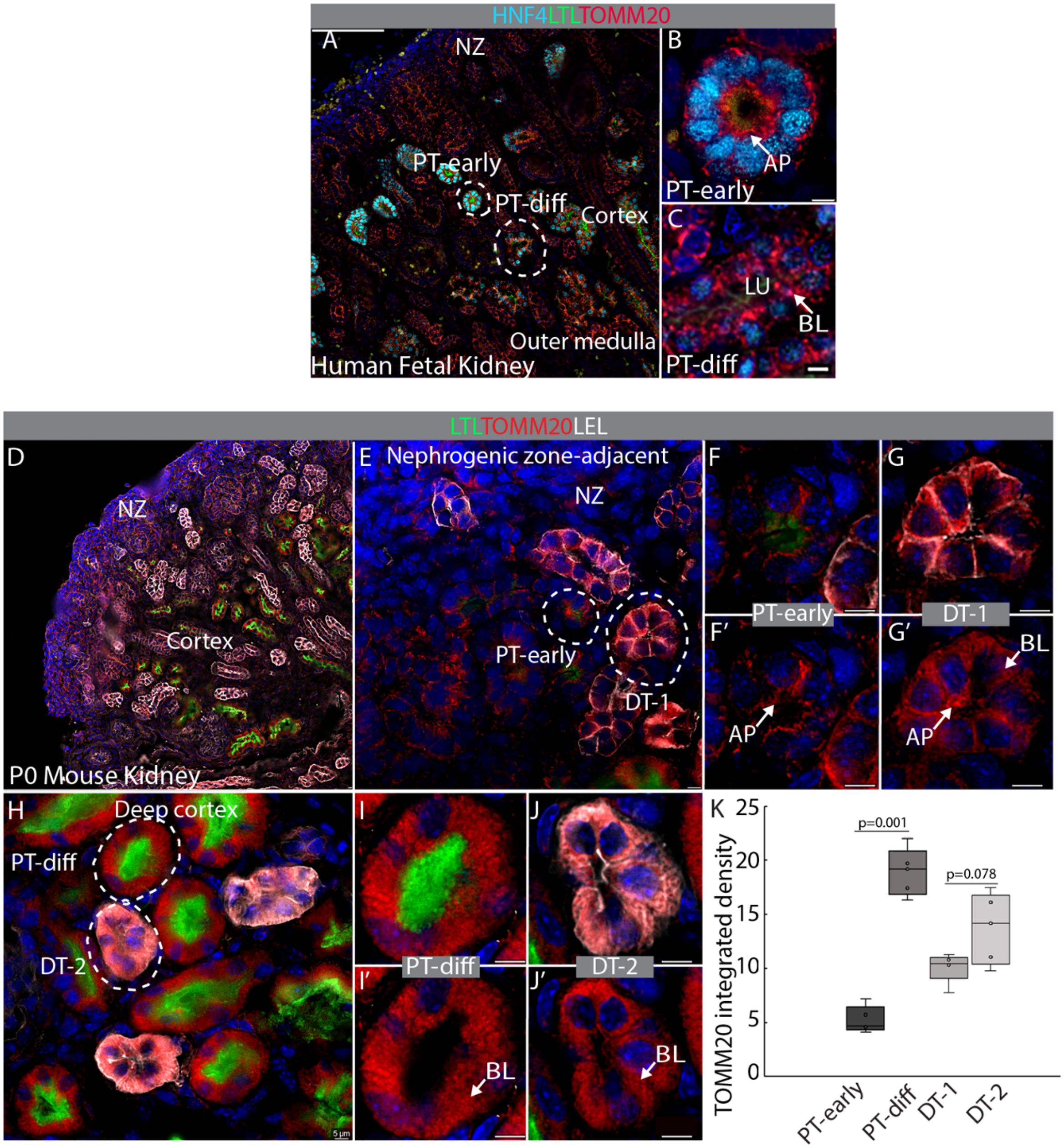


**Figure S1: Supplementary data for Figure 1.** (A-C) Week 18-20 human fetal kidney stained with HNF4A (PTs, cyan); LTL (PTs, green) and TOMM20 (mitochondria, red) (A) Widefield image showing nephrogenic zone, cortex and medulla and location of nephrogenic zone-adjacent HNF4A$^+$/LTL$^+$ PTs (PT-early) with narrow lumens and deep cortical HNF4A$^+$/LTL$^+$ PTs (PT-diff) with distended lumens. (B) PT-early with mainly apical mitochondria. (C) PT-diff with distended lumen and basolateral mitochondria. (D-J') P0 mouse kidney tissue section stained with LTL (PTs, green); TOMM20 (mitochondria, red) and LEL (DT, white). (D) Widefield image showing nephrogenic zone, cortex and medulla. (E) Nephrogenic zone-adjacent region showing LTL$^+$ PT-early and adjacent LEL$^+$ DT-1. (F-F') PT-early with largely apical mitochondria. (G-G') LEL$^+$ DT-1 with basolateral mitochondria. (H) Deep cortex showing LTL$^+$ PT-diff and adjacent LEL$^+$ DT-2 with basolateral mitochondria. (I-I') LTL$^+$ PT-diff and (J-J') LEL$^+$ DT-2 with basolateral mitochondria. (K) Comparison of integrated density of TOMM20/tubule in PT-early (n=4) vs PT-diff (n=4) and DT-1 (n=4) vs DT-2 (n=4). Data is shown as box-and-whisker with mean (middle line) and minimum–maximum values (whiskers); one way ANOVA with post-hoc Tukey HSD test. **Abbreviations:** AP, apical; BL, basolateral; DT-1, nephrogenic zone-adjacent distal tubule; DT-2, Deep cortical distal tubule; LU, lumen; NZ, nephrogenic zone; PT, proximal tubule. All immunofluorescence images are counterstained with the nuclear stain DAPI (blue). Scale bars in A 100 µm, all other panels 5µm.

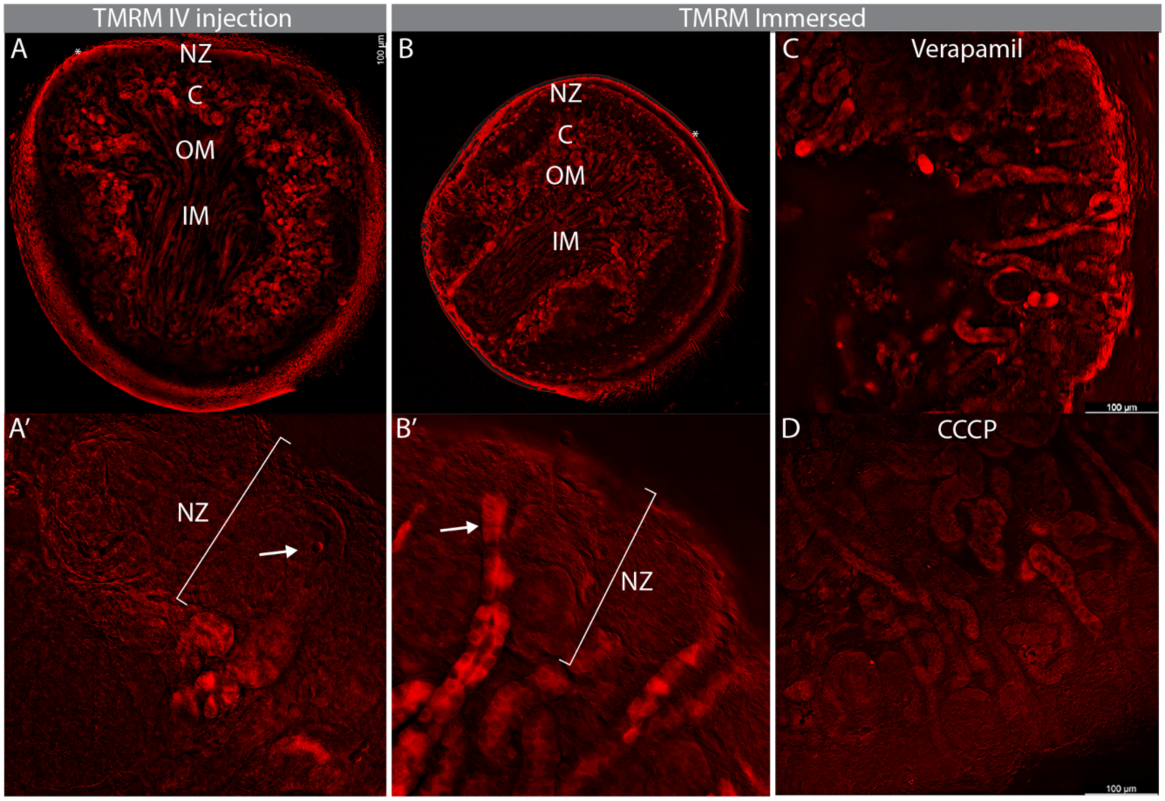


**Figure S2: Supplementary data for Figure 2.** (A) P0 kidney bisected transversely and imaged following intravenous injection of TMRM. (B) P0 kidney bisected, immersed in TMRM, briefly rinsed and imaged. Asterisk in A and B denotes incidental fluorescence from dye outside the tissue. Comparison of TMRM-injected (A') versus TMRM-immersed (B') kidneys shows that signal in nephrogenic zone (NZ) is limited to immersed kidneys (arrows). (C) Kidney treated with verapamil (MDR pump inhibitor) prior to and during immersion in TMRM. (D) Kidney treated with CCCP (mitochondrial membrane potential depolarizer) prior to immersion in TMRM to visualize fluorescence background.
**Abbreviations:** C, cortex; IM, inner medulla, OM, outer medulla, NZ, nephrogenic zone.

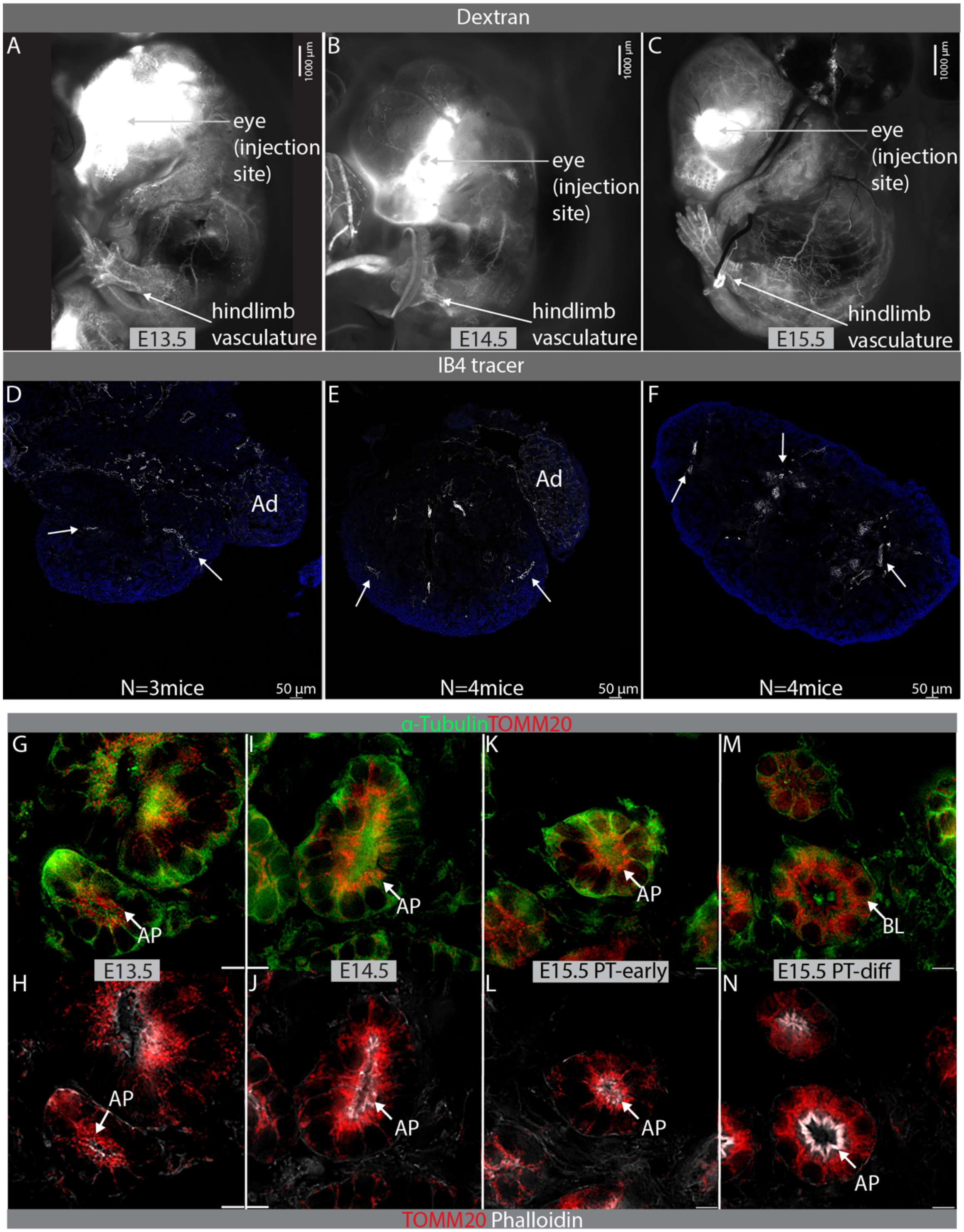

Dextran
A
B
C
1000 µm
eye
(injection
site)
hindlimb
vasculature
E13.5
E14.5
E15.5
IB4 tracer
D
E
F
Ad
N=3mice
N=4mice
50 µm
α-Tubulin TOMM20
G
I
K
M
AP
BL
E13.5
E14.5
E15.5 PT-early
E15.5 PT-diff
H
J
L
N
TOMM20 Phalloidin

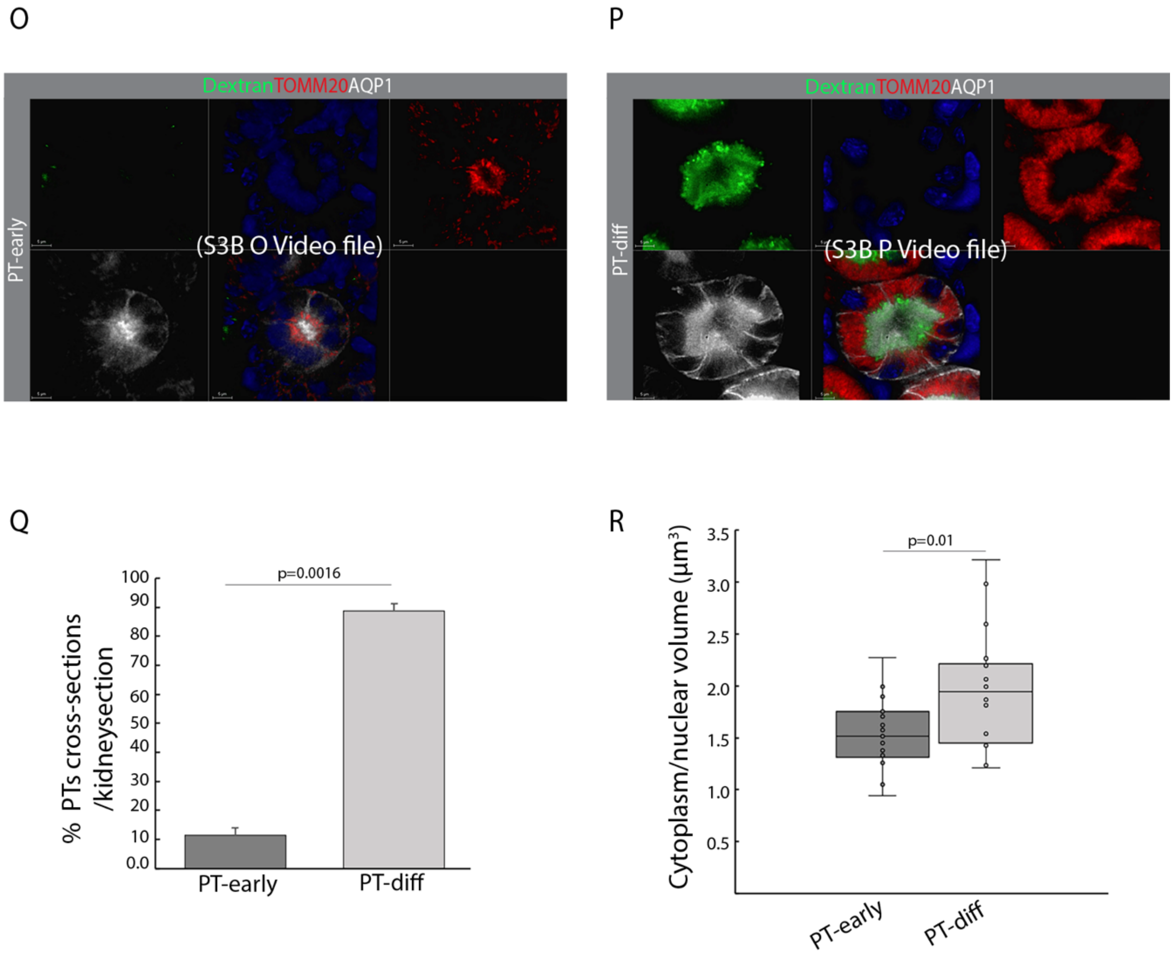


**Figure S3: Supplementary data for Figure 3.** Mouse embryos injected retro orbitally with a cocktail of Alexafluor-488-labeled 10 kD dextran and Alexafluor-647-labeled IB4. (A-C) Dextran (white) imaged 1 minute after injection to confirm dye distribution through the caudal vasculature. (D-F) Cryosections of kidneys harvested 30 minutes after injection showing IB4 binding to kidney vasculature (arrows), demonstrating distribution of the dye cocktail to the kidney. (G-N) PTs from E13.5 to E15.5 stained with α-tubulin (microtubules, green), TOMM20 (mitochondria, red) and, phalloidin (white) (G,H) PT-early at E13.5. (I,J) PT-early at E14.5. (K,L) PT-early at E15.5. (M,N) PT-diff at E15.5. Z-stacks of (O) PT-early and (P) PT-diff from P0 kidneys showing AQP1 (PT-white), TOMM20 (mitochondria-red), and Alexafluor-488-labeled 10 kD dextran (green). (Q) Quantification of percentage of PTs (PT-early vs PT-diff) per P0 mouse kidney (n=3 individuals). (R) Comparison of cytoplasm/nuclear volume ratio of PT-early (n=6 PTs/individual) and PT-diff (n=6 PTs/individual) from 3 individuals (P0); Data from 18 PTs/condition shown as box-and-whisker with mean (middle line) and minimum–maximum values (whiskers); two-tailed Student's *t*-test. **Abbreviations:** Ad, adrenal gland; AP, apical; BL, basolateral; PT, proximal tubule. Scale bar (A-C) 100 µm, (D-F) 50 µm, and (G-N) 5 µm.

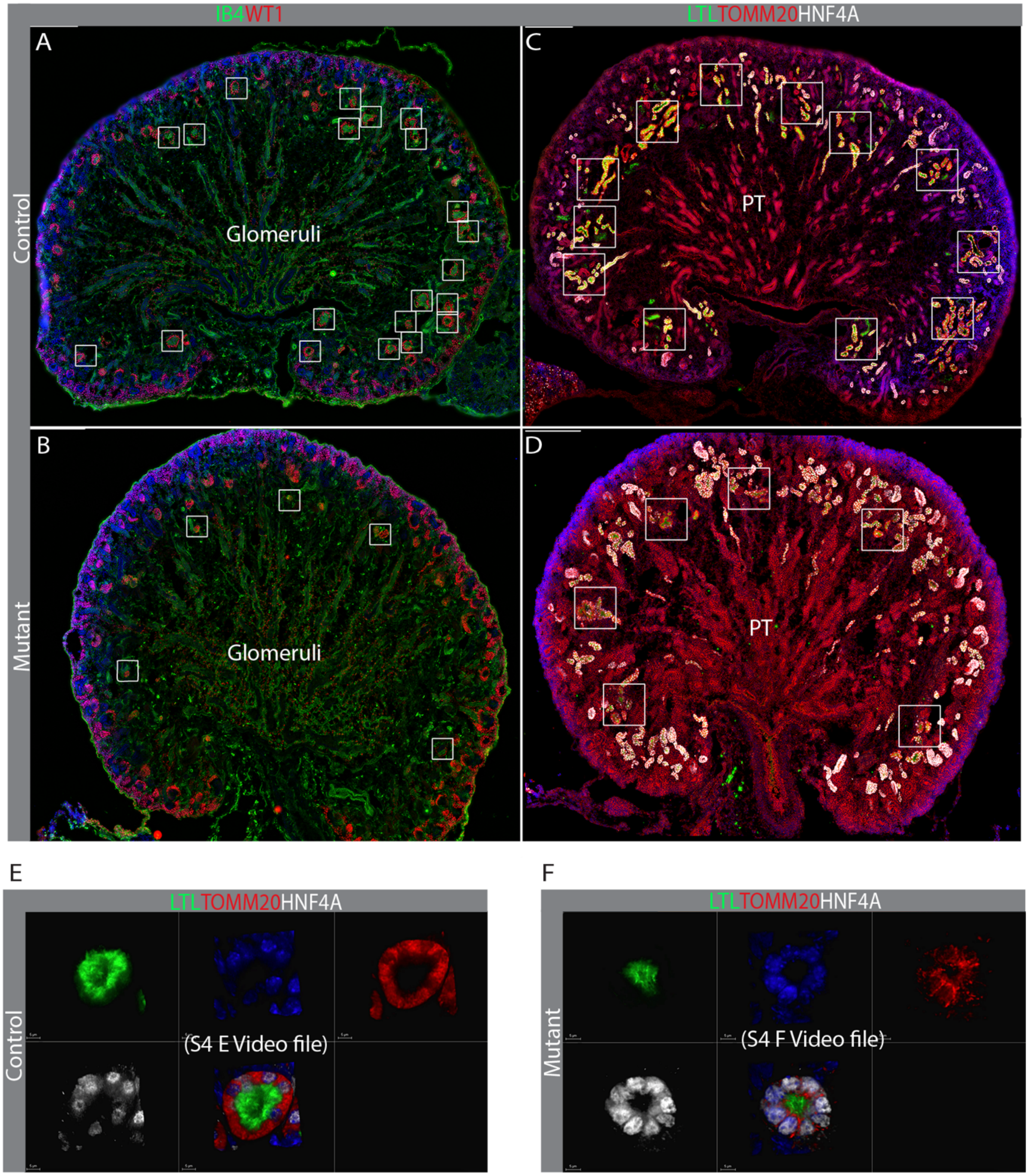
IB4WT1
LTLTOMM20HNF4A
A
C
Control
Glomeruli
PT
B
D
Mutant
Glomeruli
PT
E
F
LTLTOMM20HNF4A
LTLTOMM20HNF4A
Control
(S4 E Video file)
Mutant
(S4 F Video file)

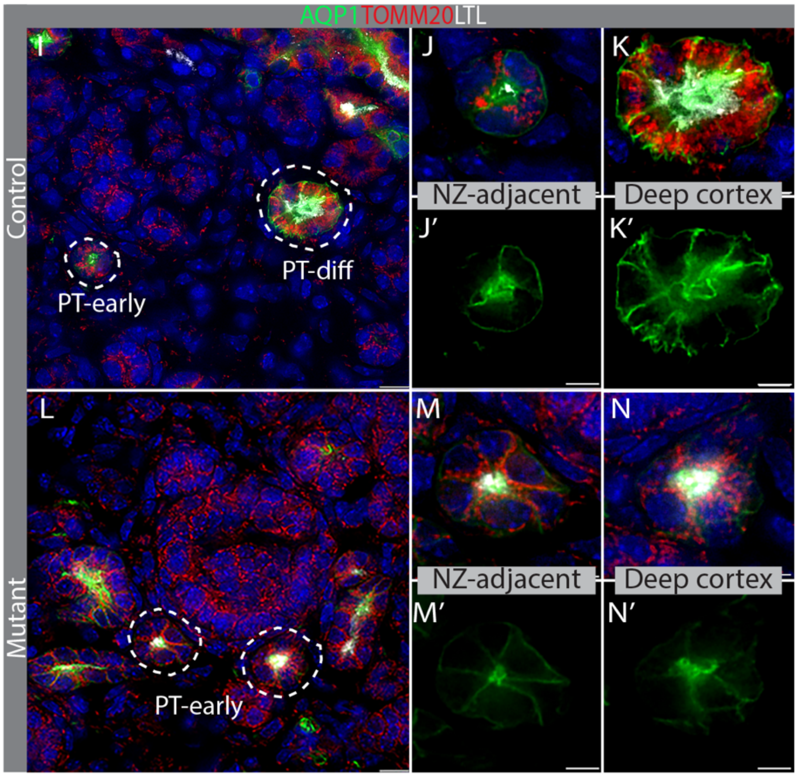


**Figure S4: Supplementary data for Figure 4.** E18.5 *Six2cre*;Numb$^{+/loxP}$;*Numbl*$^{+/loxP}$ (control) and *Six2cre*;Numb$^{loxP/loxP}$;*Numbl*$^{loxP/loxP}$ (mutant) kidneys. Representative sections of 3 wild type littermate controls. (A,B) Glomeruli identified by staining with WT1 (podocyte red) and IB4 (endothelial cell, green). (C,D) PT identified by staining for HNF4A (white) and LTL (green) and TOMM20 (mitochondria, red). Glomeruli and PT are outlined by white boxes. (A,C). (E) Maximum intensity projection and (F) Z-stack of representative control showing HNF4A/LTL (PT-white/green) and basolateral location of TOMM20 (mitochondria-red). (G) Maximum intensity projection and (H) Z-stack of representative mutant showing HNF4A/LTL (PT-white/green) and apical location of TOMM20 (mitochondria-red). (I-N') E18.5 kidneys from control and mutant stained with PT markers (AQP-1/LTL, green/white) and mitochondria (TOMM20, red). (I) Control E18.5 showing distribution of AQP1 (PT, green) in NZ-adjacent PT-early (J,J') and deep cortex PT-diff (K-K'). (L) Mutant E18.5 showing distribution of AQP1 (PT, green) in NZ-adjacent PT-early (M,M') and deep cortex PTs (N,N') which are morphologically PT-early. **Abbreviations:** PT, proximal tubule; AP, apical; BL, basolateral. All sections were counterstained with the nuclear dye DAPI (blue). Scale bars in (A-D) 200 µm, (I and L) 10 µm, (J-K') and (M-N') 5 µm.

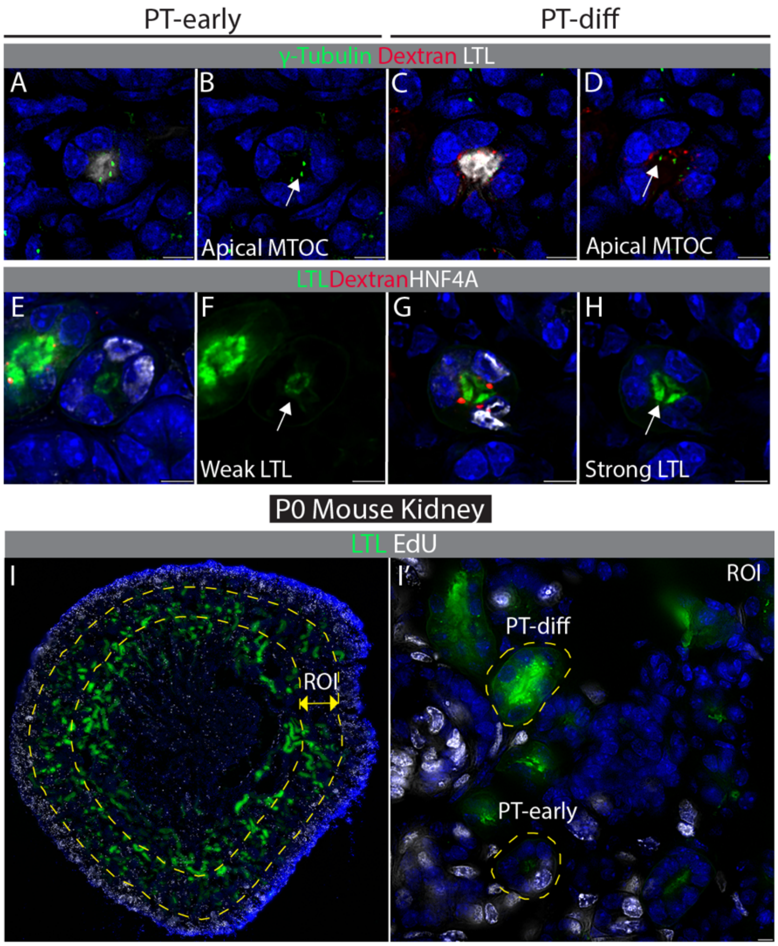


**Figure S5: Supplementary data for Figure 5.** P0 mouse kidney pulsed with 10kD dextran-Alexa568 to distinguish between PT-early (dextran negative) vs PT-diff (dextran positive). (A-D) LTL (white) and γ-Tubulin (green) to mark the microtubule organizing center. (E-H) LTL (green) and HNF4A (white). (I) and (I') LTL (PT-green) and EdU (white) to mark proliferating cells. **Abbreviations:** EdU, 5-ethynyl deoxyuridine; MTOC, microtubule organizing center; PT, proximal tubule. All sections were counterstained with the nuclear dye DAPI (blue). Scale bars in all panels 5 µm.

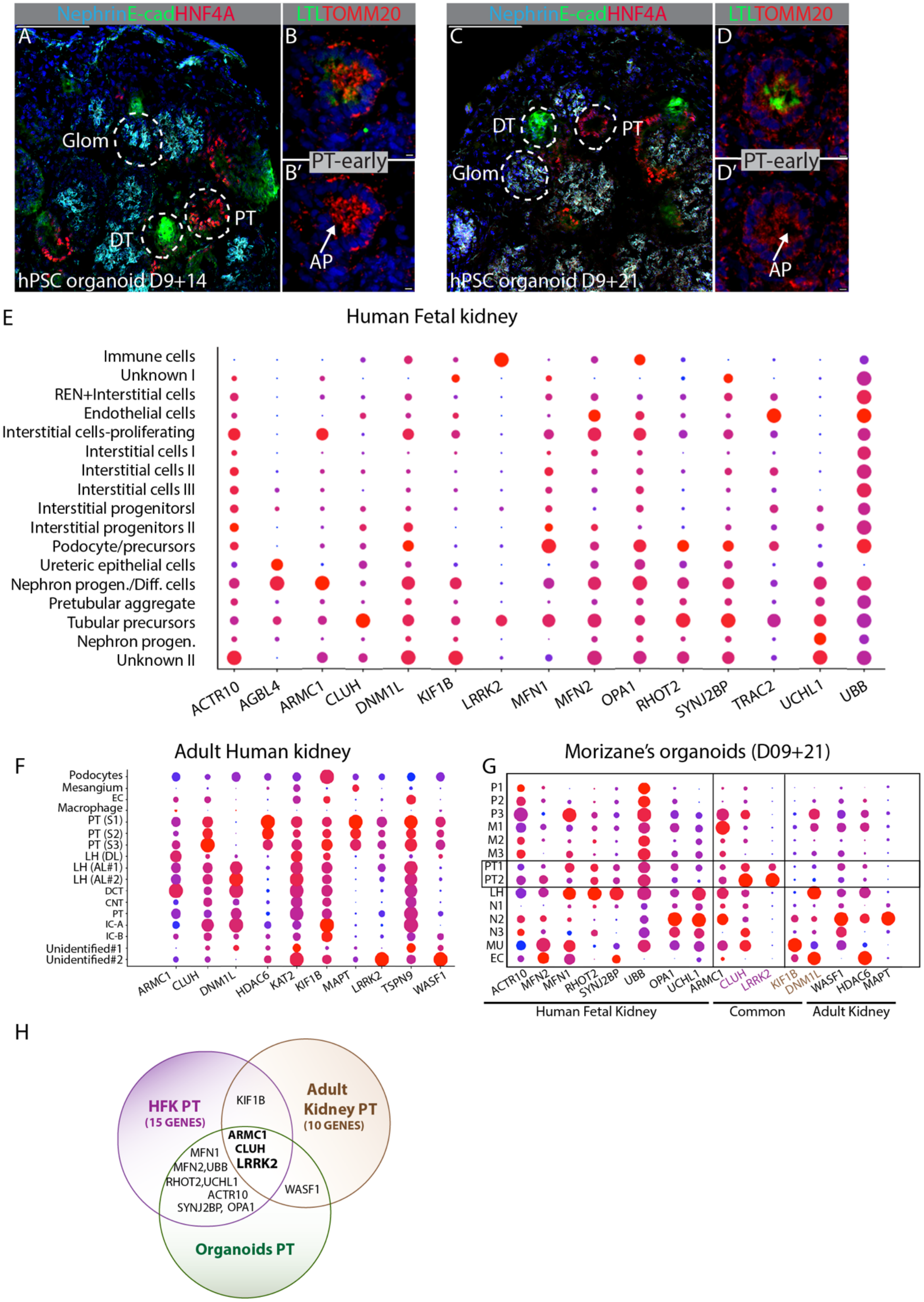

NephrinE-cadHNF4A
LTLTOMM20
A
B
C
D
Glom
PT
DT
hPSC organoid D9+14
PT-early
B'
AP
hPSC organoid D9+21
D'
E
Human Fetal kidney
Immune cells
Unknown I
REN+Interstitial cells
Endothelial cells
Interstitial cells-proliferating
Interstitial cells I
Interstitial cells II
Interstitial cells III
Interstitial progenitorsI
Interstitial progenitors II
Podocyte/precursors
Ureteric epithelial cells
Nephron progen./Diff. cells
Pretubular aggregate
Tubular precursors
Nephron progen.
Unknown II
ACTR10 AGBL4 ARMC1 CLUH DNM1L KIF1B LRRK2 MFN1 MFN2 OPA1 RHOT2 SYNJ2BP TRAC2 UCHL1 UBB
F
Adult Human kidney
Podocytes
Mesangium
EC
Macrophage
PT (S1)
PT (S2)
PT (S3)
LH (DL)
LH (AL#1)
LH (AL#2)
DCT
CNT
PT
IC-A
IC-B
Unidentified#1
Unidentified#2
ARMC1 CLUH DNM1L HDAC6 KAT2 KIF1B MAPT LRRK2 TSPN9 WASF1
G
Morizane's organoids (D09+21)
P1 P2 P3 M1 M2 M3 PT1 PT2 LH N1 N2 N3 MU EC
ACTR10 MFN2 MFN1 RHOT2 SYNJ2BP UBB OPA1 UCHL1 ARMC1 CLUH LRRK2 KIF1B DNM1L WASF1 HDAC6 MAPT
Human Fetal Kidney
Common
Adult Kidney
H
HFK PT
(15 GENES)
Adult Kidney PT
(10 GENES)
KIF1B
ARMC1
CLUH
LRRK2
MFN1
MFN2,UBB
RHOT2,UCHL1
ACTR10
SYNJ2BP, OPA1
WASF1
Organoids PT

**Figure S6: Supplementary data for Figure 6.** HNF4A (PT, red); E-Cadherin (distal tubules, green); nephrin (podocytes, cyan) in (A) Kidney organoids differentiated from H9 hESCs at 23 days (9+14, 9 days differentiation in adherent cell culture followed by 14 days non-adherent culture of aggregated cells). (B,B') PT labeled with LTL (green) and TOMM20 (red), (B') PT labeled with TOMM20 alone. HNF4A (PT-red); E-Cadherin (distal tubules-green); nephrin (podocytes-cyan) in (C) Kidney organoid differentiated from H9 hESCs at 9+21 days of culture. (D) Overlay of PT labeled with LTL (green) and TOMM20 (red), and (D') TOMM20 alone. (E-G) Expression analysis of genes in GO term G0051646 "mitochondrion localization" from single cell datasets rendered using Kidney Interactive Transcriptomics (http://humphreyslab.com/SingleCell/) in (E) Human fetal kidney, (F) Human adult kidney and (G) iPSC-derived kidney organoids annotated for genes expressed in the human fetal dataset, the adult dataset, and genes common to all 3 datasets (Common). (H) Venn diagram displaying overlap in expression of G0051646 genes between the 3 datasets. **Abbreviations**: AP, apical; DT, distal tubule; pGlom, presumptive glomerulus; PT, proximal tubule. All immunofluorescence images counterstained with the nuclear stain DAPI (blue). Scale bar in A and C 100 μM, all other panels 5 μM.

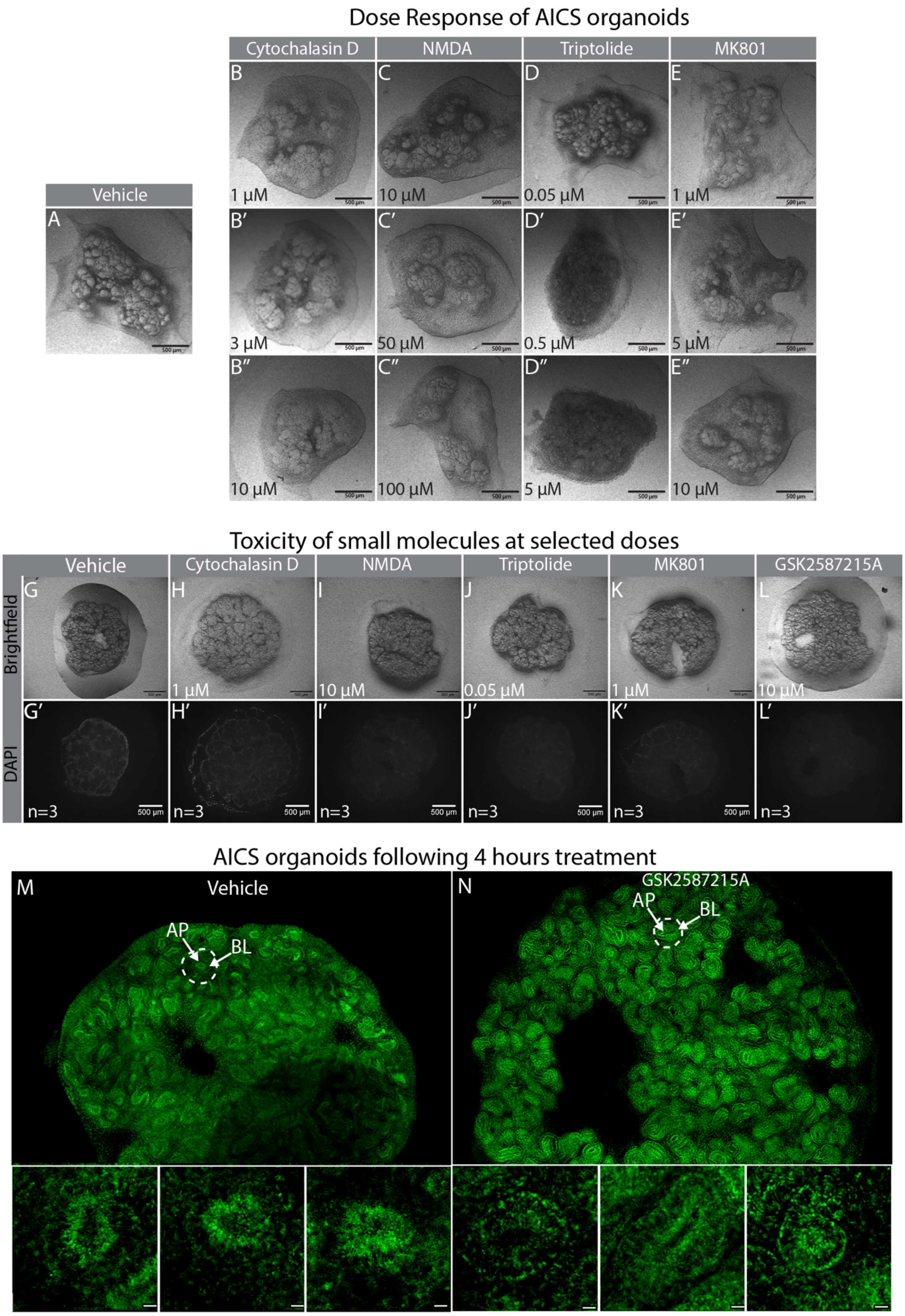
Dose Response of AICS organoids
Vehicle
A
Cytochalasin D
NMDA
Triptolide
MK801
B
C
D
E
1 µM
10 µM
0.05 µM
1 µM
B′
C′
D′
E′
3 µM
50 µM
0.5 µM
5 µM
B″
C″
D″
E″
10 µM
100 µM
5 µM
10 µM
Toxicity of small molecules at selected doses
Brightfield
DAPI
Vehicle
Cytochalasin D
NMDA
Triptolide
MK801
GSK2587215A
G
H
I
J
K
L
1 µM
10 µM
0.05 µM
1 µM
10 µM
G′
H′
I′
J′
K′
L′
n=3
n=3
n=3
n=3
n=3
n=3
500 µm
AICS organoids following 4 hours treatment
M
Vehicle
AP
BL
N
GSK2587215A
AP
BL

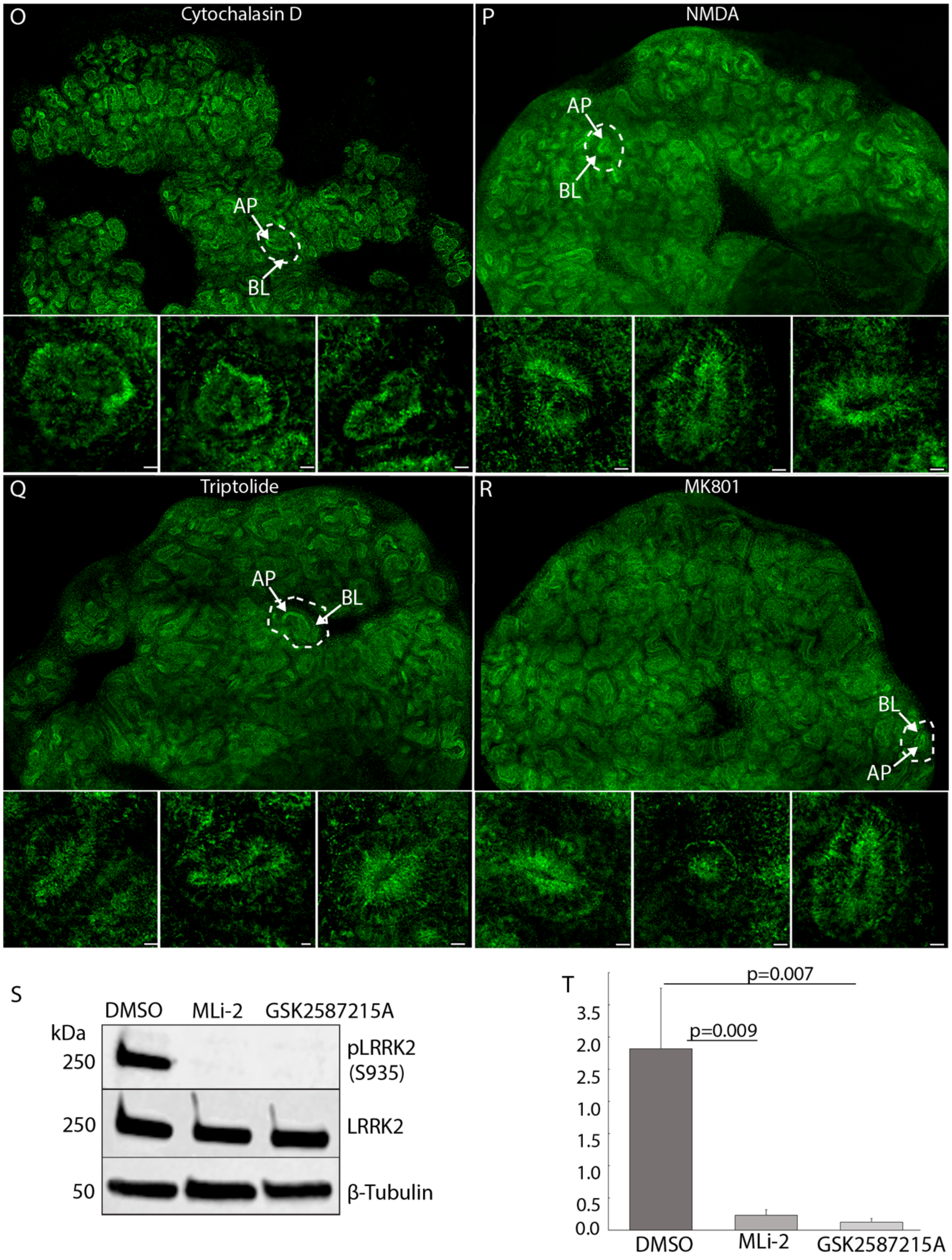
O
Cytochalasin D
AP
BL
P
NMDA
AP
BL
Q
Triptolide
AP
BL
R
MK801
BL
AP
S
DMSO
MLi-2
GSK2587215A
kDa
250
pLRRK2
(S935)
250
LRRK2
50
β-Tubulin
T
p=0.007
p=0.009
3.5
2.0
2.5
1.0
1.5
1.0
0.5
0.0
DMSO
MLi-2
GSK2587215A

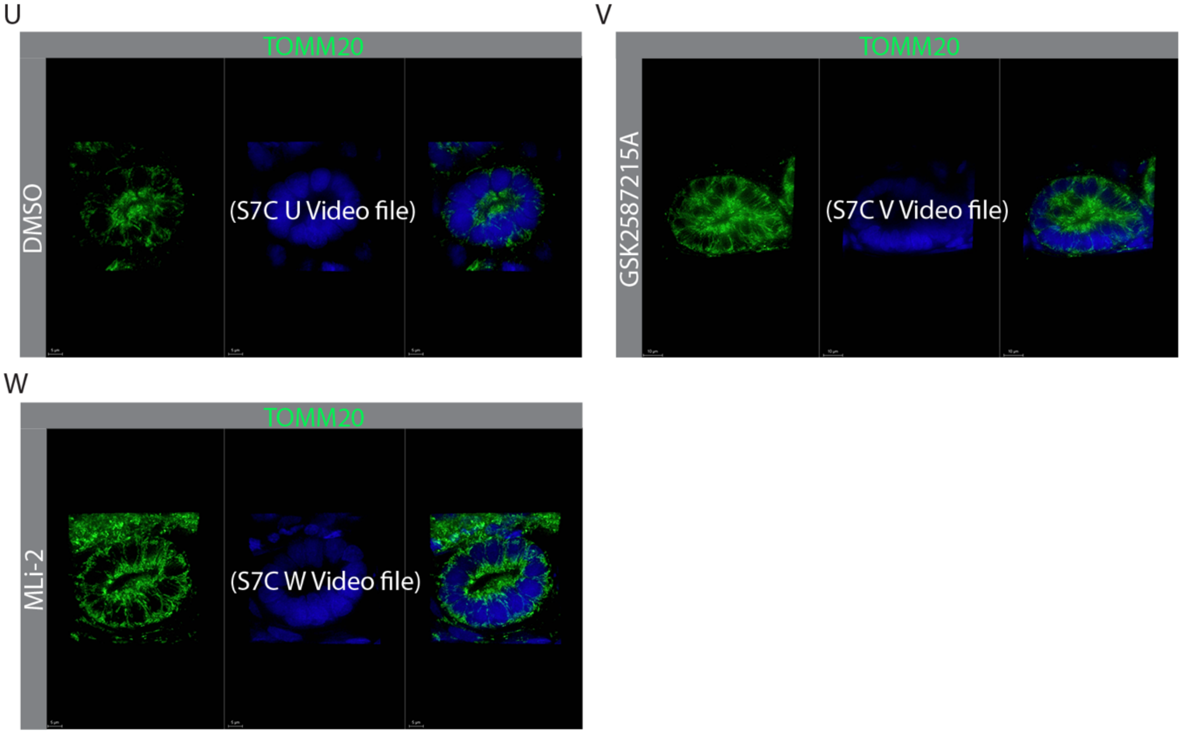


**Figure S7: Supplementary data for Figure 7.** (A-E”) Titration of compound dose on AICS-0078-79-derived kidney organoids (D9+11); representative images of 3 organoids per dose are shown. (A) DMSO, (B-B”) cytochalasin D, (C-C”) NMDA, (D-D”) Triptolide, (E-E”) MK801. (G-L’) Determination of toxicity of small molecules on AICS-0078-79 iPSCs derived kidney organoids (D9+14) by visualizing uptake of DAPI as a measure of cell death. (G-G’) DMSO (n=3), (H-H’) cytochalasin D (n=3), (I-I’) NMDA (n=3), (J-J’) Triptolide (n=3), (K-K’) MK801(n=3), (L-L’) GSK2587215A (n=3). (M-R) Whole mount images of representative AICS-0078-79 iPSCs derived kidney organoids (D9+14) showing mitochondria (green). 3 insets per organoid show examples of close-ups of individual tubules that were used for quantification of apical versus basolateral fluorescence intensity. Treatments: (M) DMSO (n=3), (N) GSK2587215A (n=3). (O) cytochalasin D (n=3), (P) NMDA (n=3), (Q) Triptolide (n=3), (R) MK801(n=3). (S) Lysates from AICS-0078-79 iPSCs derived kidney organoids (n=5) treated with vehicle or LRRK2 inhibitors immunoblotted for the active phosphorylated form of LRRK2 (pLRRK2), total LRRK2 (LRRK2) or loading control (β-tubulin) (n=3). (T) Densitometric quantification of immunoblot (n=3) is plotted as bar graph; two-tailed Student’s *t*-test. (U-W) 3D movie of AICS-0078-79 iPSCs derived kidney organoids (D9+14) mitochondria (green) counterstained with DAPI, treated with (U) DMSO (n=3), (V) GSK2587215A (n=3), (W) MLi-2 (n=3). **Abbreviations:** AP, apical; BL, basolateral. Scale bar (A-L’) 500 μM and (M-R) 100 μM.

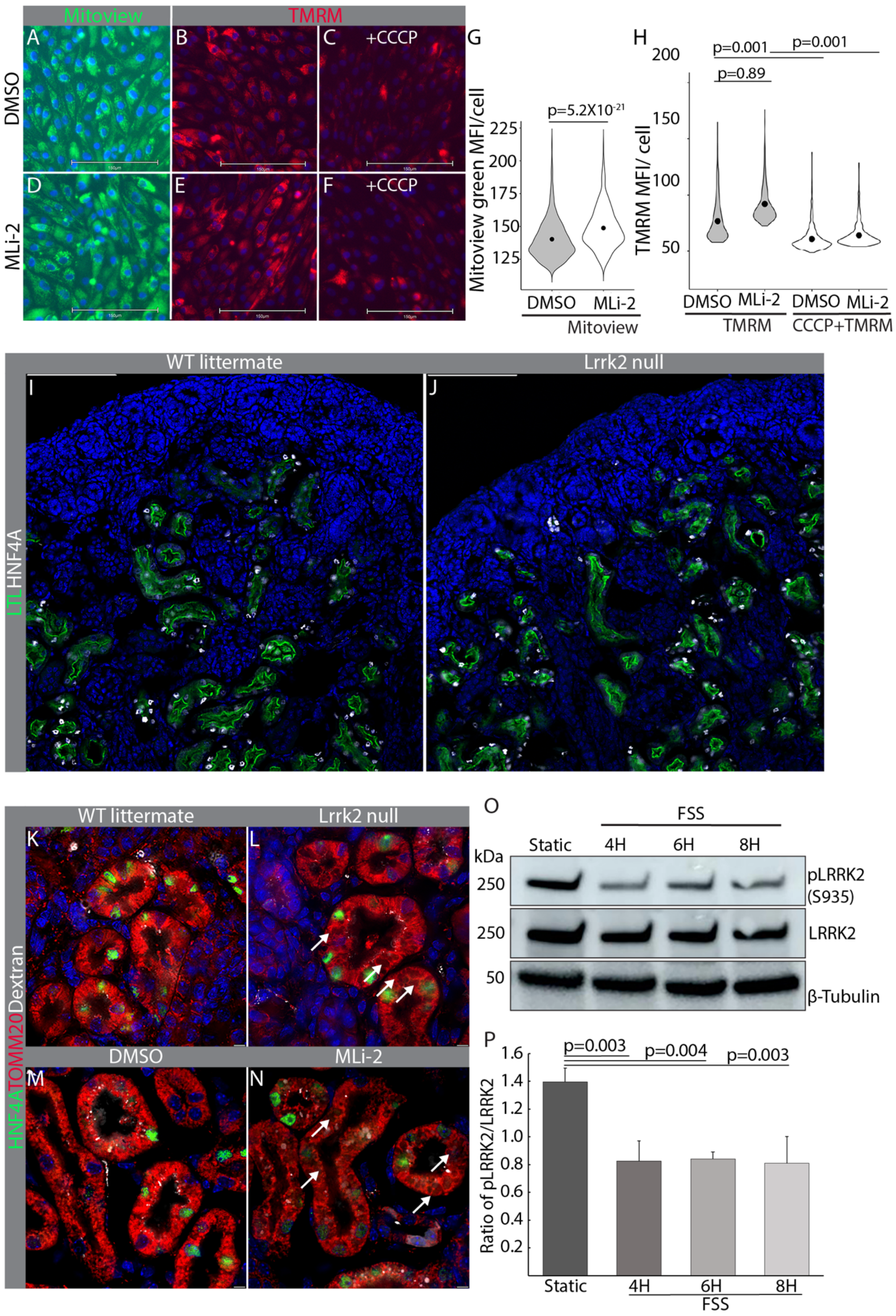
Mitoview
TMRM
A
B
C +CCCP
D
E
F +CCCP
DMSO
MLi-2
G
p=5.2X10-21
Mitoview green MFI/cell
225
200
175
150
125
DMSO
MLi-2
Mitoview
H
200
p=0.001
p=0.001
p=0.89
150
100
50
TMRM MFI/ cell
DMSO
MLi-2
DMSO
MLi-2
TMRM
CCCP+TMRM
WT littermate
Lrrk2 null
I
J
LTLHNF4A
WT littermate
Lrrk2 null
K
L
DMSO
MLi-2
M
N
HNF4ATOMM20Dextran
O
FSS
Static
4H
6H
8H
kDa
250
pLRRK2 (S935)
250
LRRK2
50
β-Tubulin
P
p=0.003
p=0.004
p=0.003
1.6
1.4
1.2
1.0
0.8
0.6
0.4
0.2
Ratio of pLRRK2/LRRK2
Static
4H
6H
8H
FSS

**Figure S8: Supplementary data for Figure 8.** hTert-RPTEC cells treated with DMSO (A-C) or MLi-2 (D-F) and stained with (A,D) Mitoview Green for total mitochondria or (B,C,E,F) TMRM (red) for mitochondrial membrane potential. (C,F) Cells pre-treated with mitochondrial depolarizer CCCP to confirm membrane potential-dependence of TMRM signal. Nuclei counterstained with Hoechst 33342 (blue). (G) Quantification of mean fluorescence intensity (MFI) per cell for Mitoview Green based on 577 cells per condition. Data is represented as violin plot; two-tailed paired Student's *t*-test. (H) Quantification of TMRM and TMRM+CCCP MFI/cell based on 484 cells per condition. Data is represented as violin plot; one way ANOVA with post-hoc Tukey HSD test. (I,J) Kidneys from representative wild type (WT) and *Lrrk2* null P0 littermates (N=3 each) stained for PT markers LTL (green) and HNF4A (white). (K,L) Representative sections of wild type and *Lrrk2* null (P0) littermates (N=3 each) pulsed with 10kD dextran-Alexa647 (white) and stained with HNF4A (green) and TOMM20 (red) to label mitochondria in PT-diff. (M,N) Representative sections of P1 wild type littermates (N=3 each) treated with vehicle (DMSO) or MLi-2 18 and 4 hours before harvest, pulsed with 10kD dextran-Alexa647 (white) and stained for HNF4A (green) and TOMM20 (red). (L,N) Arrows show areas in PTC cytoplasm of *Lrrk2* null and MLi-2 treated kidneys that are devoid of TOMM20 staining. (O) Immunoblot for phosphorylated LRRK2 (pLRRK2-S935), total LRRK2 and loading control (β-tubulin) of hTert-RPTEC cultured in static condition or subjected to fluid shear stress (FSS) for 6 hours. (P) Densitometry of immunoblot (n=3 replicates) plotted as bar graph; one way ANOVA with post-hoc Tukey HSD test. Scale bars in (A-F) 5 μm, and (I-N) 150 μm.
